# The immune landscape of tumor-associated macrophage reprogramming

**DOI:** 10.1101/2024.09.25.614962

**Authors:** Florent Duval, Joao Lourenco, Mehdi Hicham, Gaël Boivin, Alan Guichard, Celine Wyser-Rmili, Nadine Fournier, Michele De Palma, Nahal Mansouri

## Abstract

Tumor-associated macrophages (TAMs) generally acquire immunosuppressive and tumor-promoting phenotypes, which may contribute to tumor resistance to immunotherapy. We previously showed that suppression of microRNA activity through genetic *Dicer1* inactivation rewires TAM’s transcriptomes and prompts their immunostimulatory activation. This phenotypic switch enhanced recruitment and activation of CD8^+^ cytotoxic T cells (CTLs) and improved the efficacy of immunotherapy in mouse cancer models. Here, we performed single-cell RNA sequencing of whole tumors grown in either wild-type mice or mice with macrophage-specific *Dicer1* deletion. The analysis of multiple cell populations, including several discrete monocyte and macrophage subsets, indicated broad and convergent immunostimulatory programming of the tumor microenvironment, which was dependent on CTL-derived interferon-gamma (IFNγ), in mice with DICER-deficient macrophages. Intriguingly, dynamic inferences on monocyte/macrophage ontogeny and differentiation by pseudotime analysis revealed trajectories associated with progression into cell cycle, monocyte-to-macrophage differentiation, and transition from an immunostimulatory to an immunosuppressive phenotype in tumors with DICER-proficient macrophages. *Dicer1* deficiency interfered with this trajectory and stalled TAMs at an intermediate state between immature monocytes and macrophages with T cell-stimulatory capacity, thereby impeding immunosuppressive TAM development. This translated into enhanced response to antiangiogenic immunotherapy in an immunotherapy-resistant model of non-small cell lung cancer. Cycling/M2-like macrophages were conserved in human melanoma and hepatocellular carcinoma and should represent a more promising therapeutic target than the bulk of TAMs.

## Introduction

Macrophages residing in distinct tissue microenvironments display different phenotypes and functions ^1^. Such heterogeneity is defined, in part, by the identity of the circulating monocytic precursor or tissue-resident macrophage precursor from which the macrophage derives and the microenvironmental factors to which they are locally exposed ^2^. When engaged, different macrophage receptors activate distinct intracellular molecular pathways that prompt a variety of differentiation trajectories and activation states in the macrophages. Mirroring Th1 and Th2 activation of lymphocytes, macrophages may be either classically (M1) or alternatively (M2) activated ^3^. Classically activated macrophages respond to interferon-γ (IFNγ) and bacterial toxins to instigate tissue inflammation, whereas alternatively activated macrophages respond to interleukin 4 (IL-4) and other Th2 cytokines to promote angiogenesis and wound healing. Classic and alternative activation represent extremes of a phenotypic continuum, and intermediate/mixed activation states largely predominate *in vivo* ^3,4^. In tumors, macrophages often acquire an alternatively activated, M2-like phenotype that is thought to enhance tumor growth and progression ^5–8^. Reprogramming tumor-associated macrophages (TAMs) toward an M1-like phenotype has been shown to limit tumor-associated immunosuppression, leading to improved anti-tumor immunity ^5,6,8^.

Multiple cytokines expressed in the tumor microenvironment, such as IL-4, IL-10, transforming growth factor-β (TGFβ), tumor necrosis factor-α (TNFα), CCL2, colony-stimulating factor 1 (CSF1), and vascular endothelial growth factor A (VEGFA), are known to regulate macrophage development, recruitment, differentiation, and/or M1/M2-like activation in cancer ^4,9^. Many of these cytokines also modulate microRNA (miRNA) expression and activity in different cell types, including cultured monocytes/macrophages ^10^. Macrophages express measurable levels of several hundred miRNA species, some of which influence macrophage development, differentiation, and activation ^10^. We and others have previously reported that mice with an inactivating mutation in the miRNA-processing enzyme, *Dicer1*, specifically in myeloid cells exhibit delayed tumor growth in multiple cancer models ^11,12^. This effect was shown to be driven by the depletion of miRNA in macrophages, which rewired prospective TAMs to an M1-like phenotype characterized by increased IFNγ/STAT1 signaling and enhanced immunostimulatory functions ^11^. M1-like TAM programming facilitated recruitment and activation of CD8^+^ T cells that, in turn, further enforced the M1-like polarization of *Dicer1*-deficient TAMs to inhibit tumor growth. Here we leveraged *Dicer1* inactivation in TAMs to study the immune landscape of M1-like macrophage reprogramming in tumors.

## Results

### Macrophage-specific *Dicer1* deletion reprograms macrophages, delays tumor growth, and increases intra-tumoral infiltration of CD8^+^ T cells

We genetically inactivated the *Dicer1* gene in macrophages using the *LysM*-Cre mouse line. Briefly, we generated DICER-deficient mice with biallelic inactivation of *Dicer1* specifically in myeloid-lineage cells (D^KO^ hereon). We then inoculated *Dicer1* wild-type (D^WT^) and D^KO^ mice with MC38 colon adenocarcinoma cells subcutaneously (s.c.), and found that D^KO^ mice exhibited delayed tumor growth as compared to D^WT^ (**Fig. 1A**), in agreement with previous studies ^11^. We then used an orthotopic model of *Kras*^G12D^;*Tp53*^null^ non-small cell lung cancer ^13^, and found significantly smaller lung nodules in D^KO^ compared to D^WT^ mice (**Fig. 1B-C**). MC38 tumors of D^KO^ mice had higher infiltration of Ly6C^+^F4/80^−^ inflammatory monocytes but lower infiltration of Ly6C^−^F4/80^+^ macrophages and, in particular, M2-like MRC1^+^ macrophages (**Suppl. Fig. 1A-B**). Moreover, both tumor models exhibited higher CD8^+^ T cell infiltration **(Fig 1D-E** and **Suppl. Fig. 1A-B**). These results illustrate that macrophage-specific *Dicer1* inactivation inhibits tumor growth, (re)polarizes macrophages toward an M1-like phenotype, and enhances CD8^+^ T cell infiltration in the tumors.

**Fig 1.**
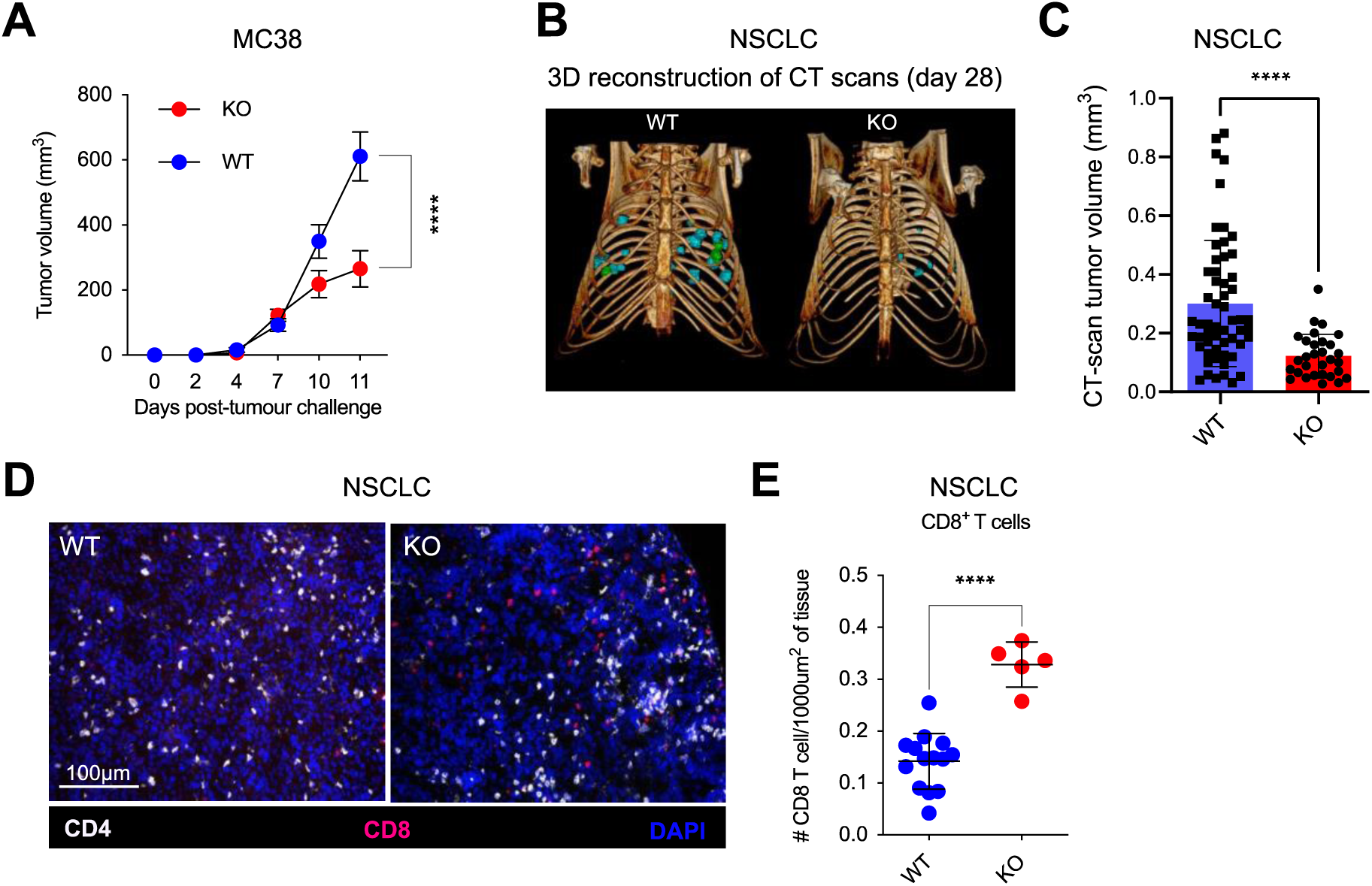
Macrophage-specific *Dicer1* deletion reprograms macrophages, delays tumor growth and increases intra-tumoral infiltration of CD8^+^ T cells. A. MC38 tumor volume in D^WT^ (*n=6*) and D^KO^ (*n=5*) mice. Statistical analysis by unpaired Student’s *t*-test. B. Representative reconstructed CT images of D^WT^ and D^KO^ mice imaged 28 days after inoculation of *Kras*^G12D^;*Tp53*^null^ cancer cells. C. Lung tumor volumes measured on CT scans 28 days post-cancer cell inoculation. Each dot represents a single tumor from D^WT^ (*n=4*) and D^KO^ (*n=5*) mice (*n*=56 total tumors in D^WT^, *n=30* in D^KO^ mice). Statistical analysis by unpaired Student’s *t*-test. D. Representative images of immunofluorescence staining of lung tumors from D^WT^ and D^KO^ mice, showing CD4^+^ (white) and CD8^+^ (red) T cell infiltration. Nuclei are stained with DAPI (blue). Scale bar, 100μm. E. Quantification of CD8^+^ T cells in lung tumors of D^WT^ (*n=4*) and D^KO^ (*n=5*) mice (*n=14* tumors analyzed for D^WT^ and *n=5* for D^KO^). Statistical analysis by unpaired Student’s *t*-test.

### D^KO^ reprograms the tumor microenvironment (TME) through IFNγ

To investigate the impact of D^KO^-induced macrophage reprogramming on the tumor microenvironment (TME), we conducted single-cell RNA sequencing (scRNA seq) of MC38 tumors obtained from either D^WT^ or D^KO^ mice at days 11-12 after MC38 tumor challenge.

We annotated different cell types using canonical marker expression (**Fig. 2A-B**, **Suppl. Fig. 2A**). We used an established signature to identify MC38 cells ^14^. The monocyte-macrophage population, characterized by high *Lyz2* and *Csf1r* expression (collectively referred to as MonoMac from now on), constituted approximately 95% of the immune cells. In line with our flow cytometric analysis (see Fig. 1E and Suppl. 1A above) CD8^+^ T cells were enriched in D^KO^ tumors, whereas there was no difference in the numbers of MonoMac cells between the two groups (**Fig. 2C-D**, **Suppl. Fig. 2B**).

**Fig 2.**
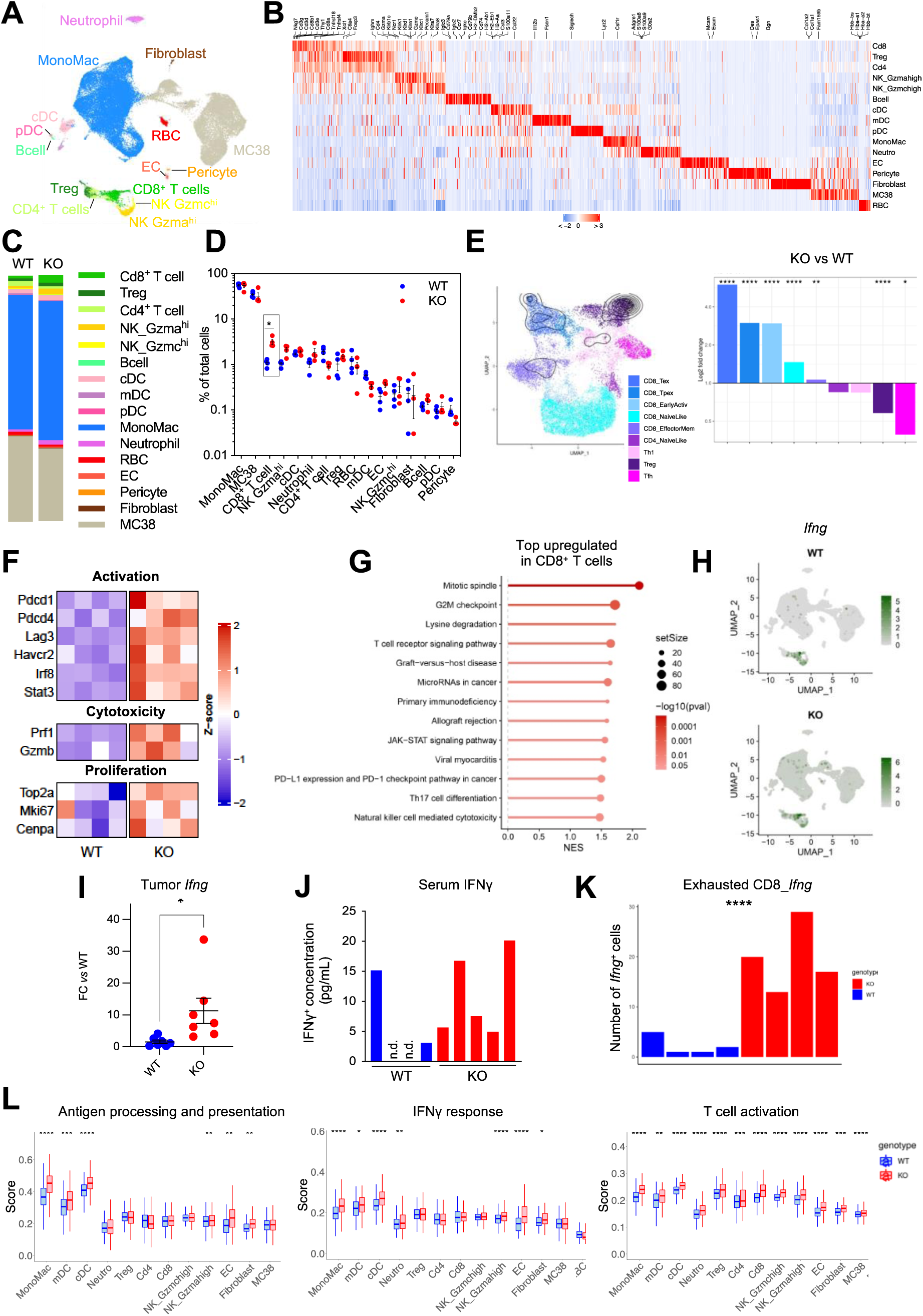
D^KO^ reprograms the microenvironment of MC38 tumors through IFNγ. A. UMAP showing cell populations identified in MC38 tumors of D^WT^ and D^KO^ mice (pooled data; *n=4* per group) by scRNA-seq. B. Expression of top enriched genes in the major cell populations of D^WT^ and D^KO^ mice (pooled data; *n=4* mice per group). Colors indicate the Z-scored gene expression. C. Bar plot showing the percentage of each cell population in D^WT^ and D^KO^ mice (*n=4*). D. Dot plot showing the percentage of each cell population in D^WT^ and D^KO^ mice (*n=4*). Statistical analysis by unpaired Student’s *t*-test. E. Projection of T cells onto the reference atlas of murine tumor-infiltrating T cell states from ProjecTILs (left), and bar plot showing the log fold-change in T cell state frequency between D^KO^ and D^WT^ mice (*n=4*). Statistical analysis by Pearson’s Chi-squared test. F. Heatmap representing the expression of the indicated genes in CD8^+^ T cells in individual D^WT^ and D^KO^ tumors (*n=4*). G. Lollipop plot showing the top upregulated Hallmark and KEGG pathways in D^KO^ CD8^+^ T cells compared to D^WT^ CD8^+^ T cells (*n=4*). H. UMAP showing the mRNA expression of the *Ifng* gene in D^WT^ and D^KO^ tumors (*n=4*). I. mRNA expression of *Ifng* in tumor lysates from D^WT^ and D^KO^ mice (*n=7*). Expression was normalized to the *Gapdh* housekeeping gene. Data are shown as fold change in D^KO^ versus D^WT^. J. IFNγ protein measured by ELISA in the serum of individual tumor-bearing D^WT^ and D^KO^ mice (*n=4*). n.d., non-detectable. K. Number of exhausted CD8^+^ T cells expressing *Ifng* in individual tumors of D^WT^ and D^KO^ mice (*n=4* for D^WT^*, n=5* for D^KO^). Statistical analysis by Student’s *t*-test. L. Hallmark gene set enrichment scores for the indicated biological processes in major cell populations of tumors of D^WT^ and D^KO^ mice (*n=4*). Boxes represent the interquartile range (IQR), horizontal lines represent the median, and whiskers represent the range within 1.5 times the IQR. Statistical analysis by Student’s *t*-test.

We next focused on tumor-infiltrating lymphocytes (TILs). To gain insights into TIL diversity upon D^KO^, T lymphocyte clusters consisting of Treg, CD4^+^ T cells and CD8^+^ T cells were segregated into substates by projecting the scRNA-seq data on a reference atlas of mutine TILs ^15^. Naïve-like, precursor exhausted, exhausted, and effector memory CD8^+^ T cells, as well as T_h_1 and Treg substates, were identifed in both D^KO^ and D^WT^ tumors. On the one hand, all activated CD8^+^ T cell sub-states (except effector memory) were increased in D^KO^ tumors; on the other, Tregs – which represent the predominant TIL subset in MC38 tumors – were significantly decreased (**Fig. 2E**, **Suppl. Fig. 2C**). Differential gene expression and pathway enrichment analysis in CD8^+^ T cells of D^KO^ versus D^WT^ tumors revealed upregulation of genes associated with activation (such as *Pdcd1*, *Lag3*, and *Irf8*) ^16,17^, cytotoxicity (*Prf1* and *Gzmb*) ^18^, and proliferation (*Mki67*, *Top2a* and *Cenpa*) ^19^ (**Fig. 2F**), as well as pathways related to T cell activation and cell cycle progression (**Fig. 2G**). Collectively, these results indicate that macrophage-specific *Dicer1* inactivation promotes intratumoral CD8^+^ T cell activation and proliferation.

IFNγ is a major effector of CD8^+^ T cell mediated anti-tumor immunity ^20^. We then analyzed the expression of *Ifng* in the tumors. *Ifng* was robustly detected in both D^WT^ and D^KO^ tumors, while *Ifna* and *Ifnb* were not detected (**Fig 2H, Suppl. Fig. 2D**). We also quantified the *Ifng* mRNA in MC38 tumor lysates using qPCR, and found that D^KO^ mice exhibited significantly higher *Ifng* expression (**Fig. 2I)**. Similarly, while the IFNγ protein was not detected in the serum of two out of four MC38-bearing D^WT^ mice, it was detected in all five D^KO^ mice (**Fig. 2J**). CD8^+^ T cells and granzyme (Gzma)^hi^ natural killer (NK) cells were the main cellular sources of *Ifng* in the tumors (**Suppl. Fig 2E**). Among CD8^+^ T cells, those displaying an exhausted phenotype highly expressed *Ifng.* While the expression of *Ifng* was similar at the single-cell level in D^WT^ and D^KO^ tumors, the number of *Ifng-*expressing, exhausted CD8^+^ T cells increased in D^KO^ tumors (**Fig. 2K, Suppl. Fig. 2F**).

To explore how D^KO^-induced IFNγ expression shapes the TME, we performed differential gene expression and pathway enrichment analyses between D^KO^ and D^WT^ tumors. Pathways regulated by interferons, such as antigen processing and presentation, IFNγ response, and T cell activation, were upregulated in D^KO^ tumors across cell subsets (**Fig. 2L, Suppl. Fig. 2G**). These results suggests that macrophage-specific *Dicer1* deficiency reprogrammed the TME through exhausted CD8^+^ T cell-secreted IFNγ.

### Monocyte and macrophages with heterogenous phenotypes reside in MC38 tumors

We next studied the effects of *Dicer1* deletion in the MonoMac subset, which are the most abundant immune cells in MC38 tumors (see figure 2C above). As their overall proportions were unchanged between D^WT^ and D^KO^ tumors, we hypothesized that MonoMac consisted of a heterogenous cell population whose dynamics and plasticity in D^KO^ versus D^WT^ tumors could differentially contribute to tumor growth. We performed unsupervised clustering of the MonoMac population and identified 9 transcriptionally distinct subclusters, represented in all samples regardless of *Dicer1* genotype (**Fig. 3A-B, Suppl. Fig. 3A-B**), which we annotated according to their gene signature. We excluded two underrepresented clusters (Macro_7 and Macro_Lyve1) from further analysis because they exhibited either a monocyte-derived dendritic cell signature (Macro_7) or a tissue-resident macrophage phenotype potentially derived from adjacent skin tissue (Macro_Lyve1). We called the first cluster Mono_1, because it exhibited an immature monocyte signature characterized by high expression of *Ly6c2*, *Plac8* and *Chil3* ^21,22^. Macro_1 and Macro_2 displayed less defined transcriptional signatures, suggesting an intermediate/transitory state between monocytes and macrophages. The Macro_3 cluster displayed an enrichment of interferon-stimulated genes, including *Rsad2* and *Ifit3* ^23^, but expressed similar genes to Mono_1. Macro_4 expressed genes associated with hypoxia, such as *Spp1*, *Hmox1* and *Mmp12* ^24,25^. Macro_5 and Macro_6, while transcriptionally related to Macro_2, exhibited upregulated cell cycle genes (*Mki67*, *Top2a,* and *Ccnb2*) ^19,26^, indicative of a proliferative state. Notably, differential abundance analysis of the MonoMac population showed lower numbers of proliferative Macro_6 in D^KO^ mice (**Fig. 3C**). We then performed differential gene expression and pathway enrichment analysis between each MonoMac subset. Unsupervised clustering of MonoMac subclusters according to Hallmark gene sets revealed 2 main groups based on their transcriptomic signatures: Inflammatory monocytes-macrophages (Mono_1, Macro_1 and Macro_3) and proliferative macrophages (Macro_5, Macro_6 and Macro_2). Proliferative macrophages displayed downregulated pro-inflammatory pathways, such as response to interferons, and upregulated cell cycle-associated biological processes. In addition, the hypoxic Macro_4 formed a third cluster (**Fig. 3D**). We then used two established signatures for M1 and M2 macrophage phenotypes derived from either *in vitro* and *in vivo* (tumor) settings ^27,28^. Inflammatory monocytes-macrophages exhibited a signature similar to M1-polarized macrophages, whereas proliferative macrophages were more M2-like (**Fig 3E** and **Suppl. Fig 3C-D**). These findings suggest the co-existence of diverse MonoMac subpopulations with potentially opposing functions in MC38 tumors. We then compared the MonoMac subclusters identified in MC38 tumors with monocyte and macrophage subsets from a mouse melanoma model ^29^. We noted similar monocyte/macrophage subsets in these different datasets (**Fig. 3F**), suggesting conservation of myeloid clusters across mouse tumors.

**Fig 3.**
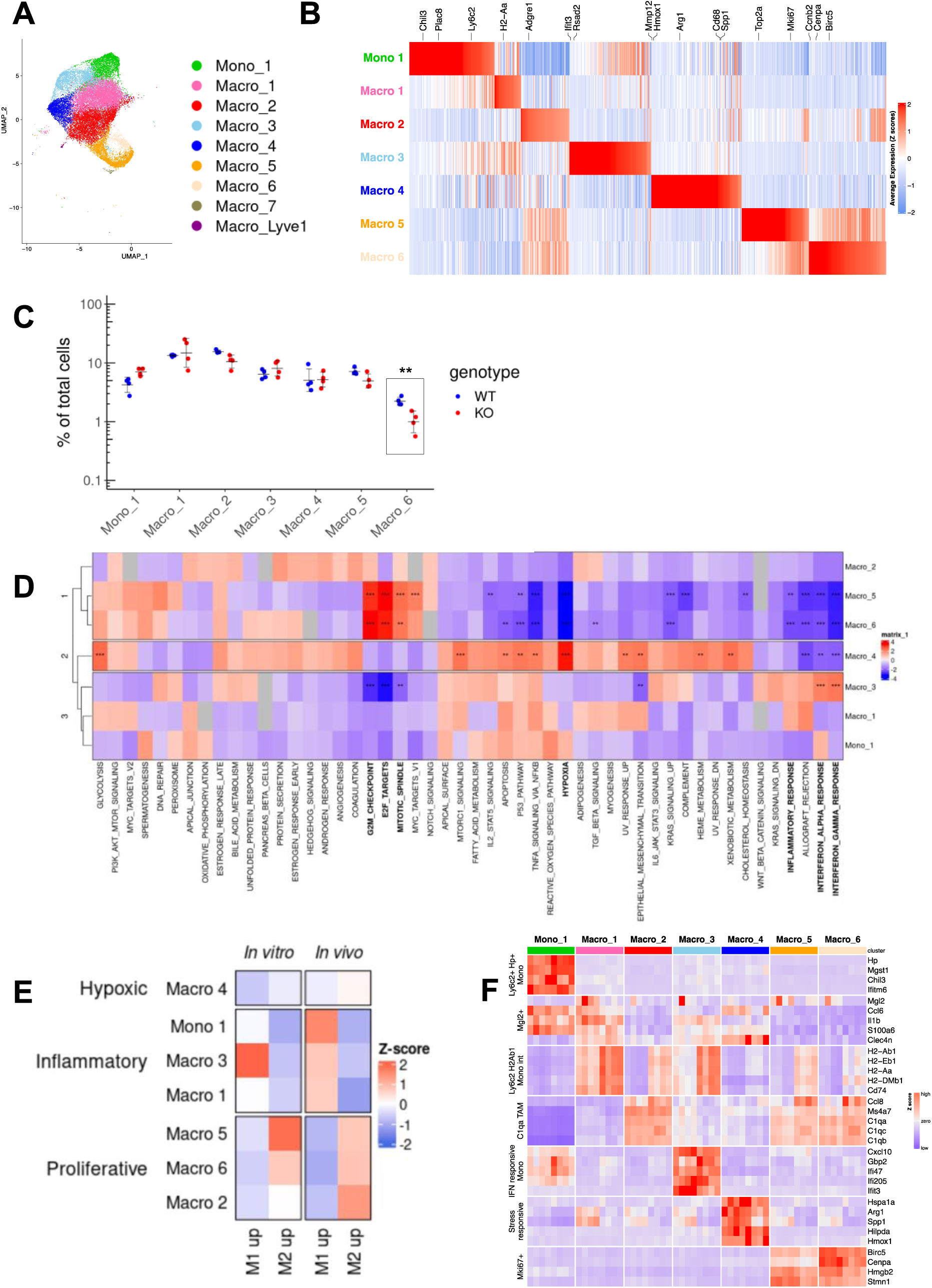
scRNA-seq reveals the co-existence of M1 and M2-like MonoMacs in MC38 tumors. A. UMAP showing subclusters within the MonoMac population of MC38 tumors of D^WT^ and D^KO^ mice (pooled data; *n=4* per group). B. Expression of top enriched genes in MonoMac subclusters of MC38 tumors of D^WT^ and D^KO^ mice (pooled data; *n=4* per group). Colors indicate the Z-scored gene expression. C. Abundance of MonoMac subclusters in D^WT^ and D^KO^ tumors (*n=4*). Bars represent the median and whiskers represent the range within 1.5 times the IQR. Statistical analysis by Student’s *t-*test. D. Normalized enrichment scores (NES) of Hallmark gene sets for each MonoMac subcluster of MC38 tumors of D^WT^ and D^KO^ mice (pooled data; *n=4* per group). Enriched genes within MonoMac subclusters were pre-ranked based on the average log2 fold-change. Colors represent the Z-scored NES. GSEA p-values adjusted for multiple testing according to Benjamini-Hochberg method. E. Signature enrichment scores for M1 and M2-like macrophage phenotypes derived from *in vitro*^28^ or *in vivo*^27^ datasets for each MonoMac subcluster of MC38 tumors of D^WT^ and D^KO^ mice (pooled data; *n=4* per group). F. Gene expression across MonoMac subclusters in individual MC38 tumors of D^WT^ and D^KO^ mice (vertical lines; *n=4*) using genes associated with “Monocyte” and “TAM” clusters from Mujal et al. 2022 (Ref ^29^).

### Dynamics analysis links trajectories and M1/M2 functionalities in MonoMacs

Concomitant presence of different MonoMac subclusters with unique or partially shared gene signatures could indicate the trajectory of differentiation within the parental MonoMac population. Macrophages present in tumors can originate from either recruitment of bone-marrow-derived monocytes and/or self-renewal of tissue-resident macrophages ^30,31^. Considering that monocyte-derived macrophages increasingly accumulate as tumors progress ^32^, we initially addressed monocyte to macrophage differentiation as a potential first trajectory. To elucidate whether the proinflammatory/M1-like and cycling/M2-like phenotypes of MonoMac are associated with monocyte to macrophage differentiation, we assessed expression of canonical markers of monocytes (*Ly6c2*) and mature macrophages (*Adgre1*, *Cd68*) in the MonoMac subsets (**Fig. 4A**). We found that monocyte markers were more highly expressed in the proinflammatory/M1-like Mono_1, Macro_1 and Macro_3 than in cycling/M2-like Macro_5, Macro_6, Macro_2. In contrast, markers of mature macrophages were higher in cycling/M2-like than in inflammatory M1-like MonoMac subclusters. Interestingly, Macro_1 and Macro_3 showed intermediate expression of canonical monocyte and macrophage markers between Mono_1 and cycling M2-like MonoMac subclusters, which could indicate a transitional state between immature monocytes and M2-like macrophages. Overall, these results suggest that MonoMac are cells distributed at different stages of monocyte to macrophage differentiation and that transition between the two is associated with a switch between M1-like to M2-like phenotypes.

**Fig 4.**
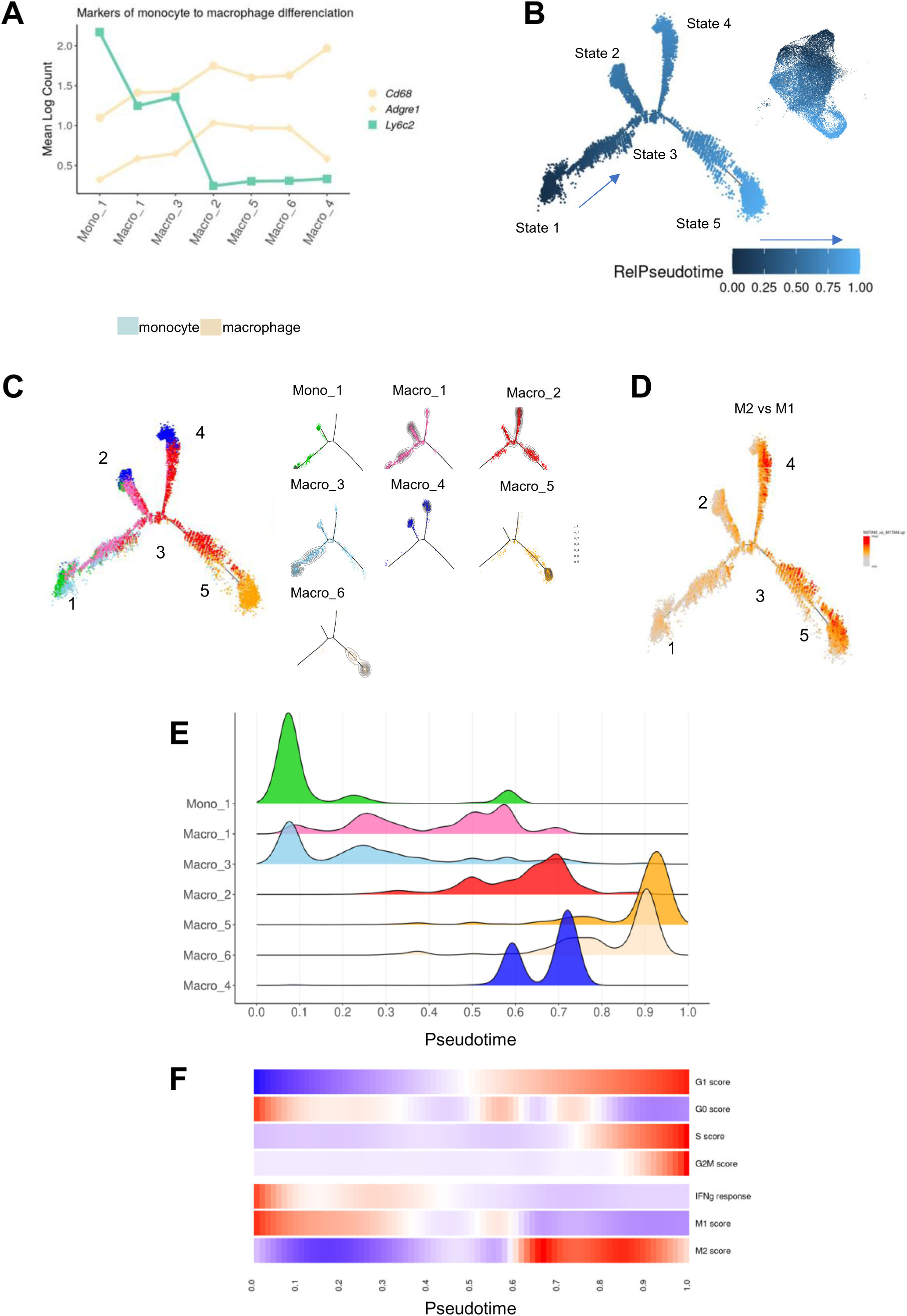
Dynamics analysis links trajectories and M1/M2 phenotypes within MonoMacs. A. Graph showing the expression of the indicated marker genes indicative of monocyte to macrophage differentiation in each MonoMac subcluster in MC38 tumors of D^WT^ and D^KO^ mice (pooled data; *n=4* per group). B. Relative pseudotime along the monocyte-macrophage differentiation trajectory, inferred by Monocle. The UMAP represents relative pseudotime in the MonoMac cluster from MC38 tumors of D^WT^ and D^KO^ mice (pooled data; *n=4* per group). The arrow indicates the direction of pseudotime. States (“branches”) in the trajectory are indicated by numbers. C. Distribution (left) and density (right) of cells from each MonoMac subcluster across the monocyte-macrophage differentiation trajectory, inferred by Monocle, in MC38 tumors of D^WT^ and D^KO^ mice (pooled data; *n=4* per group). Numbers indicate states in the trajectory. Colors indicate subclusters of MonoMac. Concentric lines indicate the level of cell density. D. Enrichment scores in MonoMac subclusters of MC38 tumors of D^WT^ and D^KO^ mice (pooled data; *n=4* per group) based on an M2-like macrophage signature derived from an *in vivo* dataset^27^, plotted along the monocyte-macrophage differentiation trajectory inferred by Monocle. Numbers indicate states in the trajectory. E. Plot representing the density of cells in each MonoMac subcluster of MC38 tumors of D^WT^ and D^KO^ mice (pooled data; *n=4* per group), relative to monocyte-macrophage differentiation pseudotime. F. Heatmap showing the enrichment of cell cycle phase, M1/M2-like macrophage phenotype, and Hallmark “interferon gamma response” gene set signatures, relative to monocyte-macrophage differentiation pseudotime. M1/M2-like macrophage phenotype signatures were derived from an *in vivo* dataset^27^. Colors indicate enrichment scores, after conditional means smoothing and zero-centering.

To interrogate trajectories within MonoMac we applied the Monocle algorithm, a method that uses pseudotime analysis to study trajectories ^33–35^. Monocle inferred five cell states stemming from state 1 (input pseudotime 0), projecting into states 2-5 through a node (state 3) (**Fig. 4B**). State 1 was mostly composed of Mono_1, Macro_1 and Macro_3, and exhibited enrichment for transcriptomic signatures of pro-inflammatory pathways (**Fig. 4C, Suppl. Fig. 4A-B**). In contrast, Macro_2, Macro_5, and Macro_6 were placed at the opposite end of the trajectory (state 5) and showed enrichment for cell cycle genes, such as E2F target and G2M checkpoint, and downregulation of the interferon signature. State 2, largely comprising Macro_1, was enriched in inflammatory pathways. State 4, largely comprising Macro_4, showed upregulation of the hypoxia pathway and downregulation of pro-inflammatory pathways. We next embedded macrophage M1/M2 signatures in Monocle trajectories. We noted a switch from M1 to M2 along the trajectory (**Fig. 4D**). This supports a trajectory of monocyte to macrophage differentiation associated with an M1 to M2 switch and the acquisition of proliferative or hypoxic phenotypes.

Since we observed that a significant fraction of M2-like MonoMac (Macro_5, Macro_6 and part of Macro_2) upregulated cell cycle genes, we speculated that the cell cycle could determine trajectories within MonoMac. To test this hypothesis, we assigned scores to each cell based on expression of genes that identify distinct cell cycle phases and plotted the scores along pseudotime obtained from Monocle ^33,36^ (**Fig. 4E-F**). Macro_2, Macro_5 and Macro_6 (state 5) aligned with active G1/S/G2M phases of the cell cycle, while an important fraction of Mono_1 and Macro_4 aligned with G0 phases. We then looked at gene expression patterns within MonoMac along the trajectory to assess whether cell cycle was associated with functional features of MonoMac. Cells within the G0/G1 phase, which corresponded to the beginning of the trajectory, upregulated IFNγ response signatures, but this pattern was lacking when the cells progressed along the cell cycle within trajectories. In line with this functional programming potentially mediated through IFNγ, we noted a gradual loss of M1-like and gain of M2-like signatures when MonoMac enter the cell cycle. We obtained similar results by applying the DeepCycle algorithm which uses RNA velocity to determine a cell cycle-driven trajectory ^37^ (**Suppl. Fig. 4C-D**). These results suggest a link between commitment to the cell cycle and monocyte differentiation into M2-like macrophages, which is paralleled with the transition from an immunostimulatory to an immunosuppressive phenotype.

### D^KO^ interferes with MonoMac trajectories and stalls TAMs in a cell cycle arrest state

Having identified that heterogeneity in the MonoMac population is associated with trajectories from a cell cycle-arrested/monocyte/M1-like to a proliferating/macrophage/M2-like state, we asked whether *Dicer1* deficiency in macrophages interferes with these trajectories. We compared D^KO^ and D^WT^ MonoMac distribution within the inferred trajectories using Monocle. We found that while the total number of MonoMac was unchanged between D^KO^ and D^WT^ (see Fig 3C above), distribution of MonoMac along trajectories was altered. Monocle showed that D^KO^ skewed MonoMacs to display a monocyte/M1-like pro-inflammatory phenotype and a cell cycle-arrested state (**Fig. 5A-D. Suppl. Fig. 5A-B**). Conversely, there were higher proliferation rates in MonoMacs progressing along trajectories and acquiring a M2-like phenotype. These results indicate that *Dicer1* inactivation interferes with MonoMac trajectories and stalls them in a more monocyte/M1-like/pro-inflammatory state, thereby restraining their maturation towards immunosuppressive M2-like macrophages.

**Fig 5.**
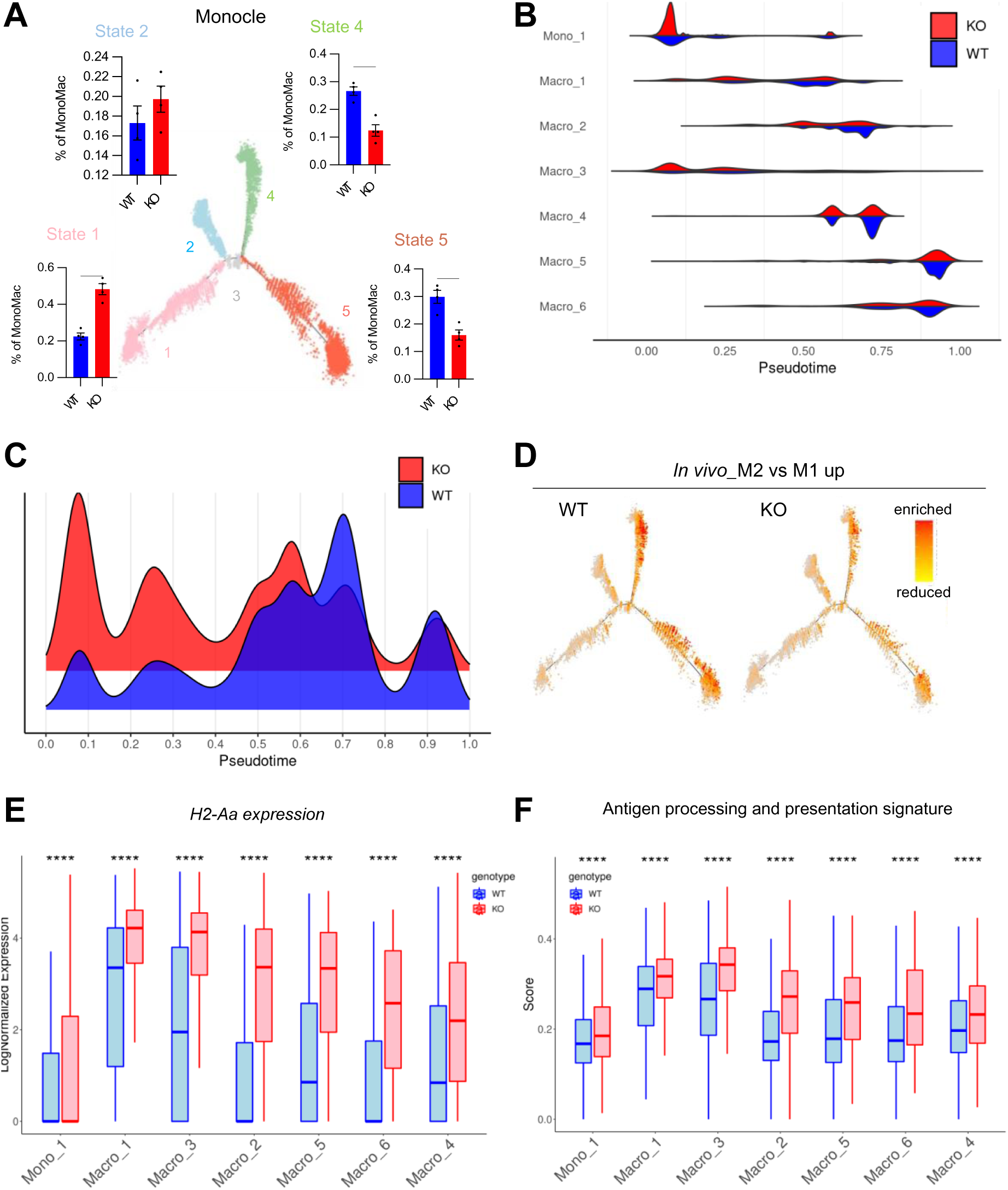
*Dicer1* inactivation in TAMs interferes with MonoMac trajectories and stalls TAMs in a cell cycle arrest state. A. Proportions of total MonoMac cells in states 1, 2, 4 and 5 of the monocyte-macrophage differentiation trajectory, inferred by Monocle by comparing tumors of D^WT^ and D^KO^ (*n=4*). B. Density of cells in each MonoMac subcluster relative to monocyte-macrophage differentiation pseudotime, by comparing tumors of D^WT^ and D^KO^ (*n=4*). C. Density of total MonoMac cells relative to monocyte-macrophage differentiation pseudotime, by comparing tumors of D^WT^ and D^KO^ (*n=4*). D. Enrichment scores for the signature of M2-like macrophage phenotypes (derived from an *in vivo* dataset^27^), by comparing tumors of D^WT^ and D^KO^ (*n=4*), plotted along the monocyte-macrophage differentiation trajectory, inferred by Monocle. E. Expression level of *H2-Aa* in each MonoMac subcluster of tumors of D^WT^ and D^KO^ mice (*n=4*). Boxes represent the IQR, horizontal lines represent the median, and whiskers represent the range within 1.5 times the IQR. Statistical analysis by Student’s *t-*test. F. Enrichment score of the “antigen processing and presentation” GOBP term gene set in each MonoMac subcluster of tumors of D^WT^ and D^KO^ mice (*n=4*). Boxes represent the IQR, horizontal lines represent the median, and whiskers represent the range within 1.5 times the IQR. Statistics using Student’s *t*-test.

We next evaluated the effect of *Dicer1* KO on antigen presentation. All MonoMac subsets upregulated the *H2Aa* gene and antigen presentation signatures in D^KO^ (**Fig. 5E-F**). We also studied *Dicer1* KO-induced interference on MonoMac trajectories using flow cytometry of the tumors. We used Ly6C and F4/80 to distinguish between immature monocytes, intermediate monocytes/macrophages, and mature macrophages, as previously described ^38,39^. We identified 3 populations of MonoMac encompassing Mono_1 (Ly6C^+^F4/80^−^), Macro_1 and Macro_3 (Ly6C^+^F4/80^+^), or Macro_2, Macro_4, Macro_5, and Macro_6 (Ly6C^−^ F4/80^+^) (**Suppl. Fig. 5C**). In agreement with scRNA-seq data, the mean fluorescence intensity (MFI) of Ly6C decreased progressively from Mono_1-like, Macro_1/3-like and Macro_2,4,5,6-like, whereas F4/80 increased. Moreover, Ly6C was upregulated in the Mono_1-like and Macro_1/3-like clusters in the D^KO^ tumors (**Suppl. Fig. 5D-E**). These findings, both at the RNA and protein levels, indicate that *Dicer1* KO affects MonoMac trajectories, causing cells to stall in a cell cycle-arrested, immunostimulatory phase.

### CD8^+^ T cell-driven IFN**γ** enriches M1-like MonoMacs in D^KO^

We have previously shown that macrophage-specific *Dicer1* deletion triggers M1-like TAM programming, marked by enhanced IFNγ/STAT1 signaling. This reprogramming abated the immunosuppressive functions of TAMs and promoted recruitment of activated CD8^+^ T cells to the tumors. In turn, CD8^+^ T cell-derived IFNγ further enhanced M1 polarization of *Dicer1*-deficient TAMs and inhibited tumor growth ^11^. Building on these findings, and since intratumoral exhausted T cells expressed highest levels of IFNγ (see Suppl. Fig. 4D above), we hypothesized that CD8^+^ T cell-derived IFNγ could be responsible of driving the perturbation of MonoMac trajectories in D^KO^ mice. To test this hypothesis, we treated MC38 tumor-bearing mice with irrelevant IgGs or an IFNγ neutralizing antibody and interrogated the distribution of Mono_1-like, Macro_1/3-like and Macro_2/4/5/6-like cells by FACS. While IgG-treated D^KO^ tumors had higher proportions of Macro_1/3-like cells and lower proportions of Macro_2/4/5/6-like cells within MonoMac, IFNγ neutralization normalized the proportions of Macro_1/3-like and Macro_2/4/5/6-like to levels similar to those in D^WT^ tumors (**Fig. 6A**). We next assessed the expression of MHCII, a marker of M1-like macrophages, and MRC1, a marker of M2-like macrophages, in each MonoMac population in D^KO^ tumors following IFNγ neutralization. While MHCII was upregulated and MRC1 downregulated in the Macro_1/3-like and Macro_2/4/5/6 cells of D^KO^ tumors, both effects were negated by IFNγ neutralization (**Fig. 6B**). These results indicate that IFNγ, likely secreted by CD8^+^ T cells, directs MonoMac trajectories.

**Fig 6.**
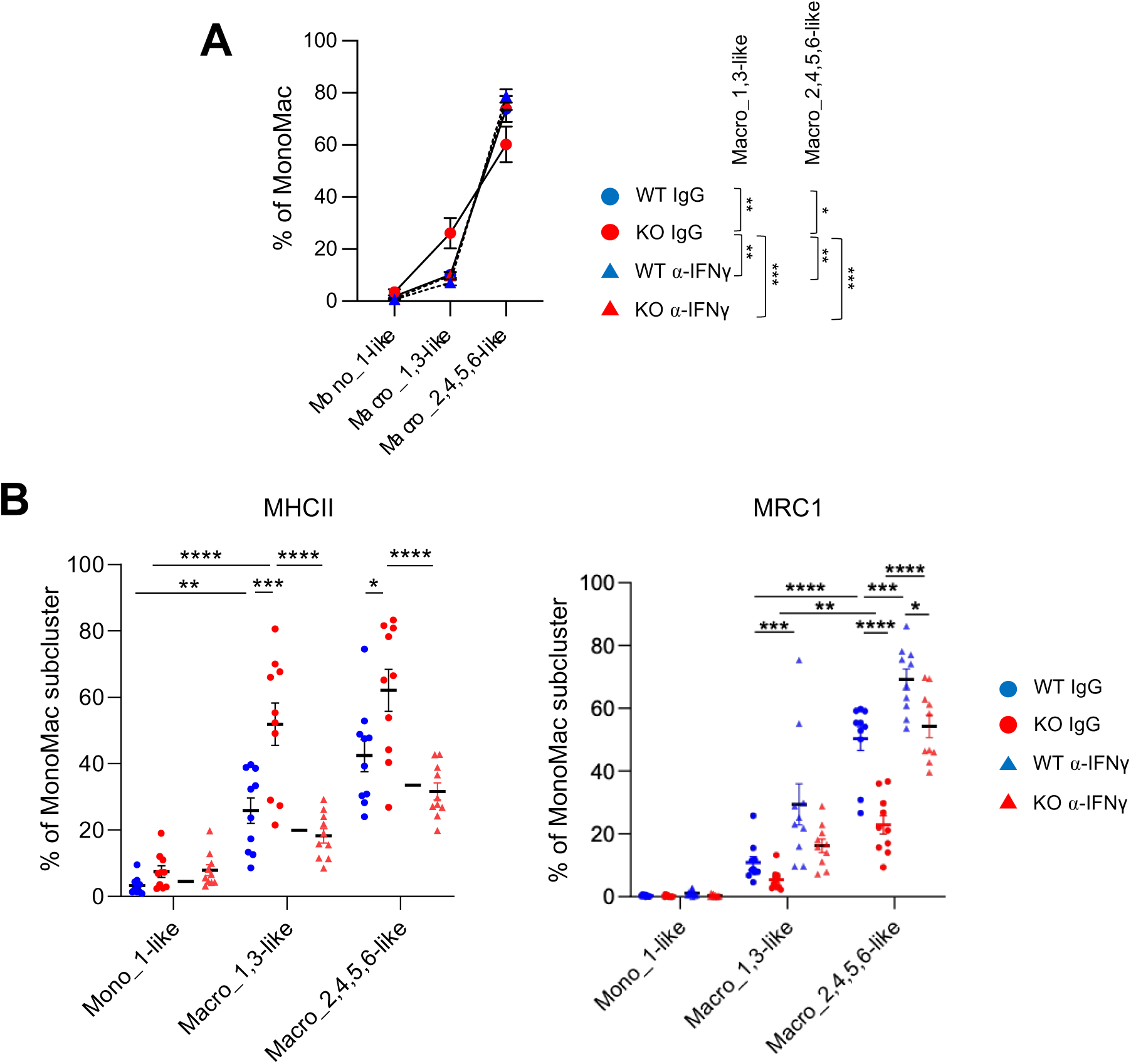
CD8^+^ T cell-derived IFNγ enriches for M1-like MonoMacs in tumors with *Dicer1* inactivation in TAMs. A. Percentage of the indicated myeloid cells in tumors of D^KO^ or D^WT^ mice (n=10) treated with an IFNγ-neutralizing antibody (α-IFNγ) or control IgGs, shown as percentage of all MonoMacs. Statistical analysis by two-way Anova. B. Percentage of MonoMacs expressing MHCII (left) or MRC1 (right) in tumors of D^KO^ or D^WT^ mice (n=10) treated with an IFNγ-neutralizing antibody (α-IFNγ) or control IgGs, quantified by FACS. Statistical analysis by two-way Anova.

### D^KO^ enhances tumor response to antiangiogenic immunotherapy

TAMs limit tumor responses to immunotherapy and its combination with other agents, such as antiangiogenic drugs ^40^, in several cancer types, including lung cancer, through immunosuppressive reprogramming of the TME ^5^. Depleting both monocyte-derived and tissue-resident TAMs enhanced tumor response to “antiangiogenic immunotherapy” (a combination of VEGFA, angiopoietin-2 and PD1 blockade) in a mouse NSCLC model ^13^. Building on these results, we evaluated whether macrophage reprogramming in D^KO^ mice could phenocopy pharmacological TAM targeting to improve tumor response to antiangiogenic immunotherapy in an orthotopic NSCLC model. To this aim, we administered A2V (a bispecific antibody against angiopoietin-2 and VEGFA ^41^) and PD1 antibodies (or their respective IgG controls) to mice 15 days post-tumor inoculation, and lung tumor growth was assessed with micro-CT. Consistent with our previous findings ^13^, combinatorial A2V and anti-PD1 treatment not only failed to inhibit tumor growth, but also led to accelerated progression of some tumor nodules in D^WT^ mice. Tumor growth was greatly inhibited in D^KO^ mice, regardless of treatment. Yet, A2V plus anti-PD1 enhanced the therapeutic effect, leading to virtually complete tumor regression in D^KO^ mice (**Fig. 7A-B**). These results demonstrate that reprogramming TAMs to a pro-inflammatory/M1-like phenotype improves the efficacy of antiangiogenic immunotherapy in an experimental lung cancer model.

**Fig. 7.**
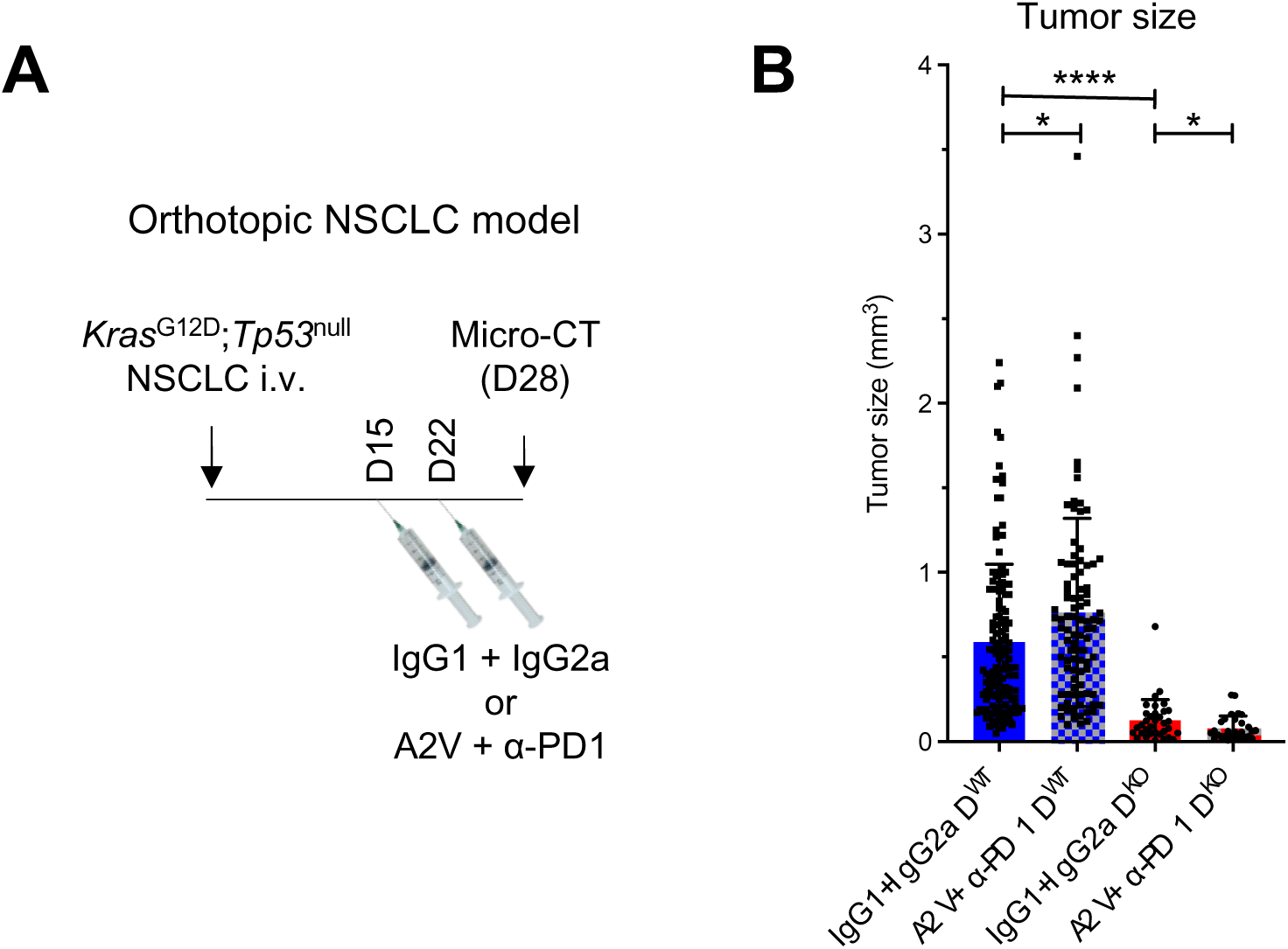
*Dicer1* inactivation in TAMs magnifies tumor response to antiangiogenic immunotherapy. A. Schematic of experiment showing the outcome of antiangiogenic immunotherapy (A2V + PD1 blockade) in an orthotopic *Kras*^G12D^;*Tp53*^null^ NSCLC model established in D^WT^ or D^KO^ mice. B. Volume of individual lung tumors in the mice treated as indicated. *n=6* D^WT^ mice treated with IgG1+IgG2a; *n=7* D^WT^ mice treated with A2V+α-PD1; *n=5* D^KO^ mice. Note that each mouse carries several independent lung tumors. Statistical analysis using unpaired Student’s *t*-test.

### The M2-like signature of proliferating macrophages is conserved in human melanoma **and hepatocellular cancer**

We investigated the presence of proliferating macrophages in publicly available scRNA-seq datasets of several cancer types ^42^. A distinct cluster of myeloid *Mki67*^+^ cells was present in melanoma and hepatocellular carcinoma (**Fig. 8A, B**). This population was highly enriched in S/G2M cell cycle signatures genes (**Fig. 8C, D**). Moreover, myeloid *MKI67*^+^ cells exhibited an M2-like transcriptional signature according to both in vitro- and in vivo-generated M1/M2 signatures (**Fig. 8E, F**). Furthermore, subdividing human monocyte and macrophage populations into either *MKI67*^+^ or *MKI67*^−^ clusters unraveled a positive correlation between G2M and M2 scores in *MKI67*^+^ monocyte/macrophages in melanoma and hepatocellular carcinoma (**Fig. 8G, H**). Together, these results show that cycling MonoMac with an M2-like phenotype are conserved in human cancers.

**Fig 8.**
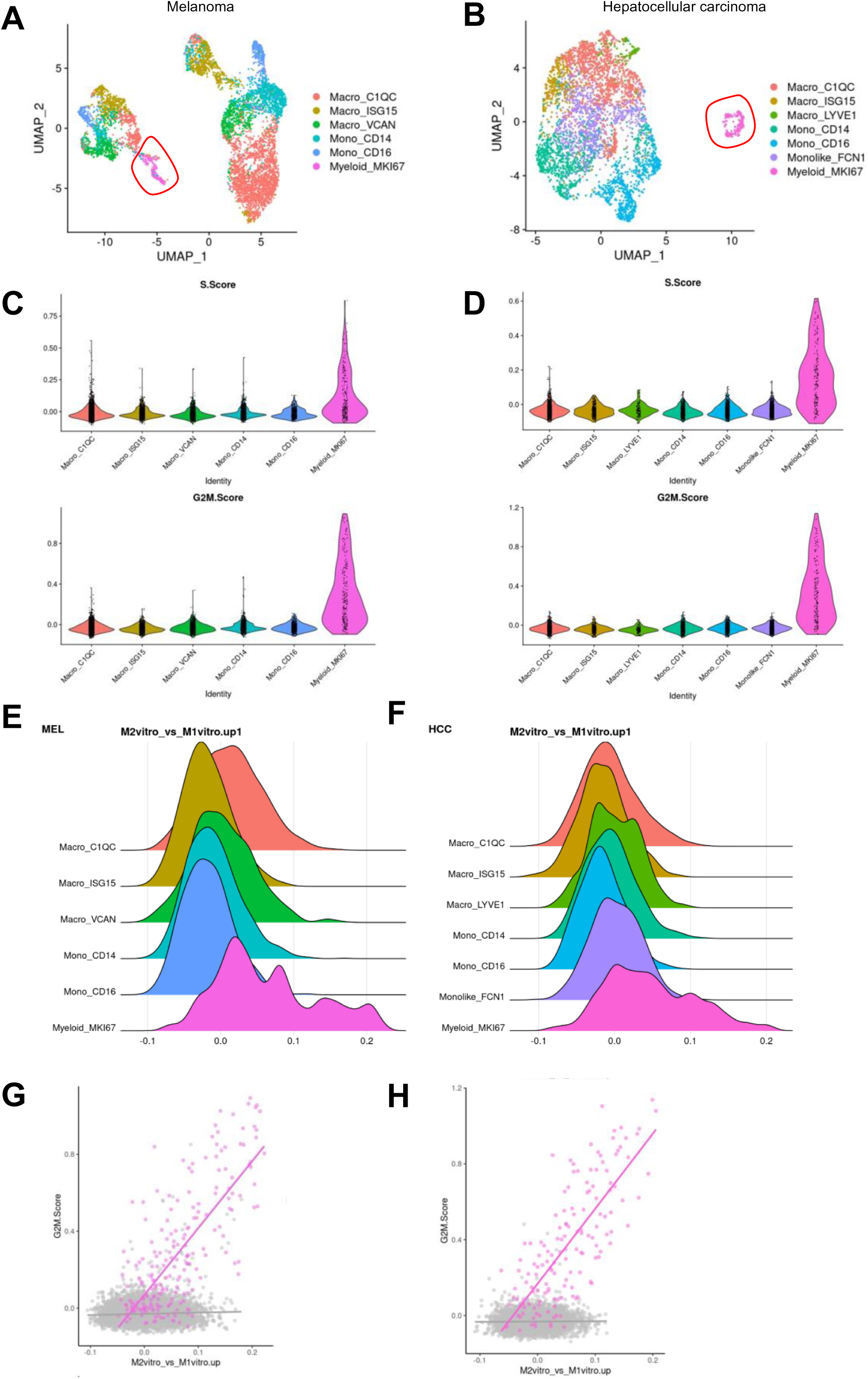
Cycling/M2-like macrophages are conserved in human cancer types. A. UMAP projection of MonoMac cells from the human melanoma (MEL) dataset. Cells are color coded by sub-population. Proliferating macrophages (“Myeloid_MKI67”) are encircled by a red line. B. UMAP projection of MonoMac cells from the human hepatocellular carcinoma (HCC) dataset. Cells are color coded by sub-population. Proliferating macrophages (“Myeloid_MKI67”) are encircled by a red line. C. Enrichment scores of the S and G2M cell cycle phase signatures across MonoMac subclusters from the human melanoma (MEL) dataset. Cells are color coded by subcluster. D. Enrichment score of the S and G2M cell cycle phase signatures across MonoMac subcluster from the human hepatocellular carcinoma (HCC) dataset. Cells are color coded by subcluster. E. Density of MonoMac subclusters from the human melanoma (MEL) dataset related to the enrichment score of the G2M cell cycle phase signature. F. Density of MonoMac subclusters from the human hepatocellular carcinoma (HCC) dataset related to the enrichment score of the G2M cell cycle phase signature. G. Correlation between the enrichment scores of the G2M cell cycle phase and M2-like macrophage phenotype signatures (derived from an *in vitro* dataset^28^) in MonoMac cells from the human melanoma (MEL) dataset (pink: proliferating macrophages, adjusted R-squared = 0.5622, p≤0.0001; gray: other cells, adjusted R-squared = 0.002709; p≤0.0001). H. Correlation between the enrichment scores of the G2M cell cycle phase and M2 macrophage phenotype signatures (derived from an *in vitro* dataset^28^) in MonoMac cells from the hepatocellular carcinoma (HCC) human dataset (pink: proliferating macrophages, adjusted R-squared = 0.5907, p≤0.0001; gray: other cells, adjusted R-squared < 0.0001; p>0.05).

## Discussion

In this study, we utilized a mouse model of macrophage-specific *Dicer1* deletion to investigate the cellular and molecular landscape of the tumor microenvironment (TME) following enforced M1-like macrophage programming. Consistent with previous findings from our group and others in murine models of primary breast cancer, colon cancer, and glioblastoma ^11,12^, we here report a tumor-protective effect associated with macrophage-specific *Dicer1* knockout (D^KO^) in both the MC38 model of colon cancer and an orthotopic model of non-small cell lung cancer (NSCLC). In both models, M1-like macrophage programming was associated with enhanced infiltration and activation of cytotoxic CD8^+^ T cells and marked inhibition of tumor growth, attesting to the pivotal role played by polarized macrophages in sculpting tumor progression.

scRNA-seq of MC38 tumor-bearing mice revealed distinct cell clusters within the Monocyte-Macrophage (MonoMac) population, regardless of *Dicer1* genotype status. These clusters were further categorized into three main subgroups: inflammatory, hypoxic, and proliferative MonoMacs. Bioinformatic analysis of potential trajectories within the MonoMac population suggested a dynamic process of monocyte differentiation toward a more terminally differentiated macrophage state, which coincides with the loss of an M1-like immunostimulatory phenotype and the acquisition of an M2-like immunosuppressive one. Further analyses indicated that these trajectories are embedded within the phases of the cell cycle. Cells at the beginning of the trajectory displayed an M1-like, cell cycle-arrested phenotype, while progression along the trajectory led to either an M2-like proliferative macrophage substate (or a quiescent hypoxic state). Of note, proliferative M2-like macrophages were observed previously in mouse mammary tumor models ^43^. Interestingly, deletion of *Dicer1* in MonoMacs skewed theses trajectories and stalled monocyte differentiation at the beginning of the trajectory in the cell-cycle arrested state associated with a M1-like, immunostimulatory state. In addition, we showed that macrophage-like MonoMacs (Ly6C^−^F4/80^+^) exhibit higher MRC1 and lower MHCII expression (typical of M2-like, immunosuppressive TAMs) in D^WT^ mice, whereas acquisition of this phenotype was mitigated in D^KO^ mice. Our findings align with a recent report, which described the loss of proliferative capacity of TAMs but gain of anti-tumoral properties upon macrophage-specific *Dicer* deletion in a mouse model of glioblastoma ^12^. Importantly, our results indicate that M1-like macrophage skewing in D^KO^ mice enhanced tumor response to a combination of antiangiogenic and immune therapy (antiangiogenic immunotherapy ^40^) in a treatment-resistant model of experimental NSCLC ^13^.

IFNγ, which is highly expressed in exhausted CD8^+^ T cells, emerged as a pivotal orchestrator of immunostimulatory programming of the TME. Unsupervised clustering of genes and pathways that were significantly deregulated in tumors of D^KO^ mice revealed upregulation of several interferon-associated genes and pathways, such as antigen processing and presentation, T cell activation, and response to interferon. Furthermore, we observed the upregulation of IFNγ mRNA and protein levels in tumors and serum of D^KO^ mice, respectively. Interestingly, pharmacological IFNγ neutralization distrupted monocyte to macrophage differentiation, irrespective of the *Dicer* genotype status, underscoring the impact of IFNγ in driving macrophage polarization towards an M1-like phenotype. This reprogramming demonstrably enhanced the immunostimulatory capacity of macrophages, thereby reinforcing the cytotoxic activities of CD8^+^ T cells.

We found that cycling MonoMacs with an M2-like phenotype were conserved in human melanoma and hepatocellular cancers. This finding should encourage investigating strategies for selectively targeting (or reprogramming) this cycling TAM subset. Of note, we previously reported that cycling TAMs in a transgenic NSCLC model were associated with tumor resistance to antiangiogenic immunotherapy; elimination of these proliferative TAM with cisplatin dramatically improved tumor response by removing a cellular source of immunosuppressive factors that sustained regulatory T cells ^13^. Because cisplatin does not specifically target immunosuppressive and cycling TAMs, alternative strategies should be explored for selectively targeting these pro-tumoral cells while sparing (or even enhancing) immunostimulatory M1-like TAMs, for example, through macrophage-targeted DICER inhibitors ^44^.

## Acknowledgements

This study was supported by grants from the ISREC Foundation (Tandem to M.D.P. and N.M.), the Swiss National Science Foundation (SNSF 310030-188868 to M.D.P., and CRSK-3_190441 to N.M.), and the European Research Council (ERC CoG EVOLVE-725051 to M.D.P.). The authors thank the Brain & Mind Institute (EPFL) for sc-RNA-Seq and acknowledge the excellent technical support of the other scientific platforms at EPFL and Agora.

## Author contributions

F.D. performed experiments, analyzed and interpreted the data, and drafted the manuscript. J.L. conducted and N.F. supervised bioinformatics analyses. M.H., G.B., A.G. and C.W.-R. contributed to the execution of some experiments and data analysis. N.M. co-supervised the study, analyzed and interpreted the data, and drafted the manuscript. M.D.P. supervised and coordinated the study, interpreted the data, and edited the manuscript.

## Supplemantary figures

**Suppl. Fig 1.**
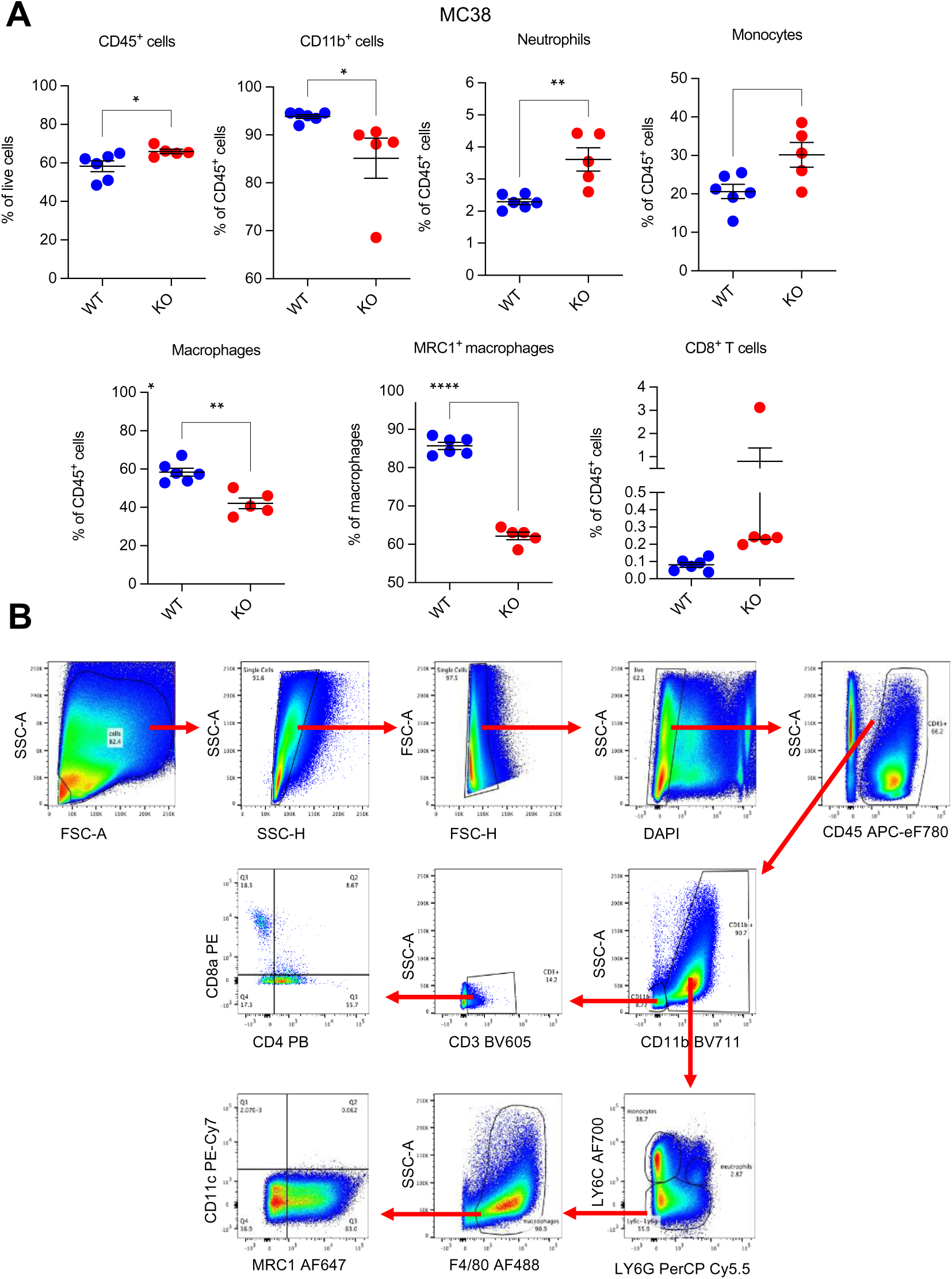
Macrophage-specific *Dicer1* deletion reprograms macrophages, delays tumor growth and increases intra-tumoral infiltration of CD8^+^ T cells. A. Percentage of the indicated cell types in MC38 tumors of in D^WT^ (*n=6*) and D^KO^ (*n=5*) mice, measured by flow cytometry. Statistical analysis by unpaired Student’s *t*-test. B. Gating strategy used for flow cytometric analysis of immune cells in MC38 tumors.

**Suppl. Fig 2.**
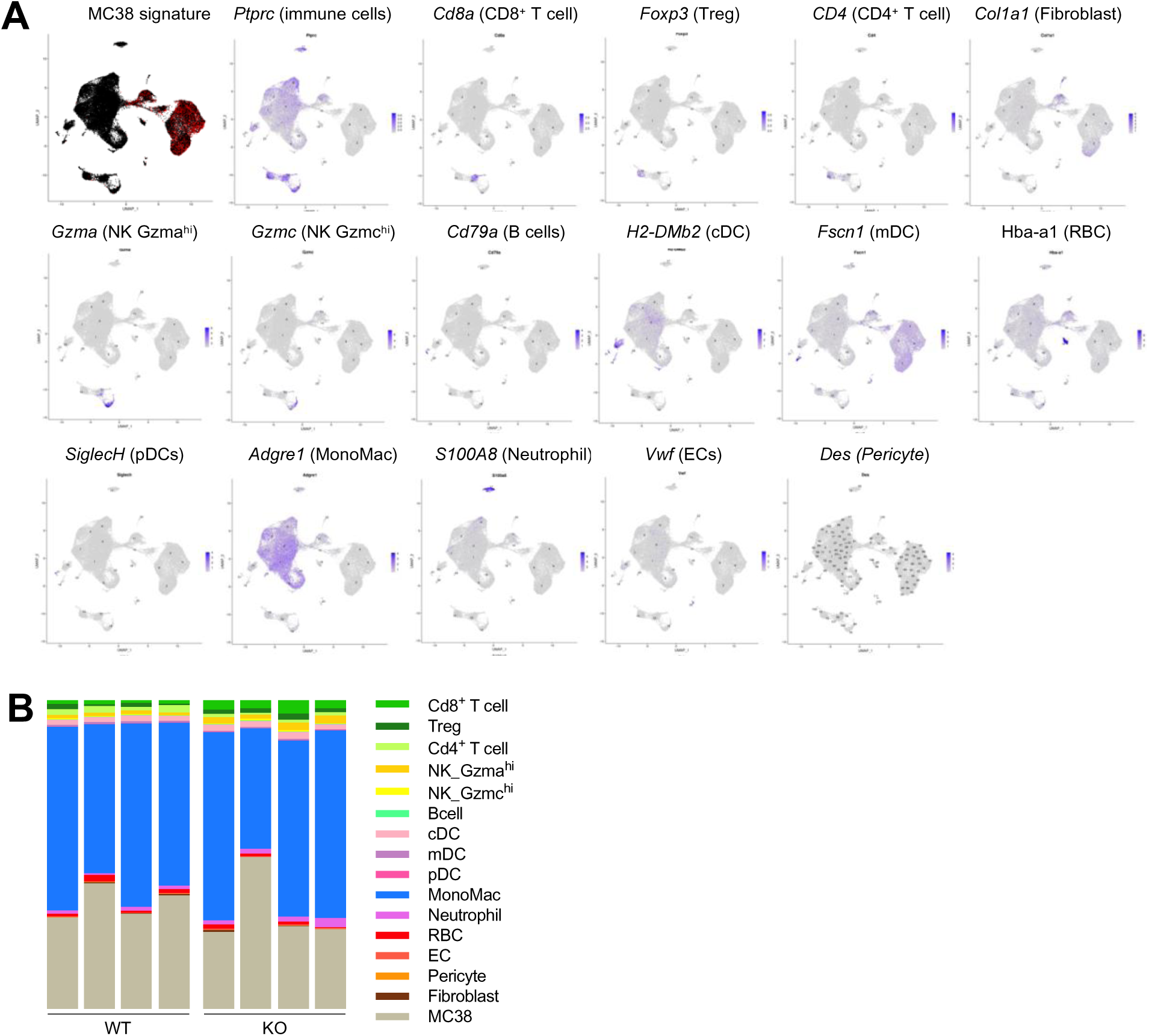
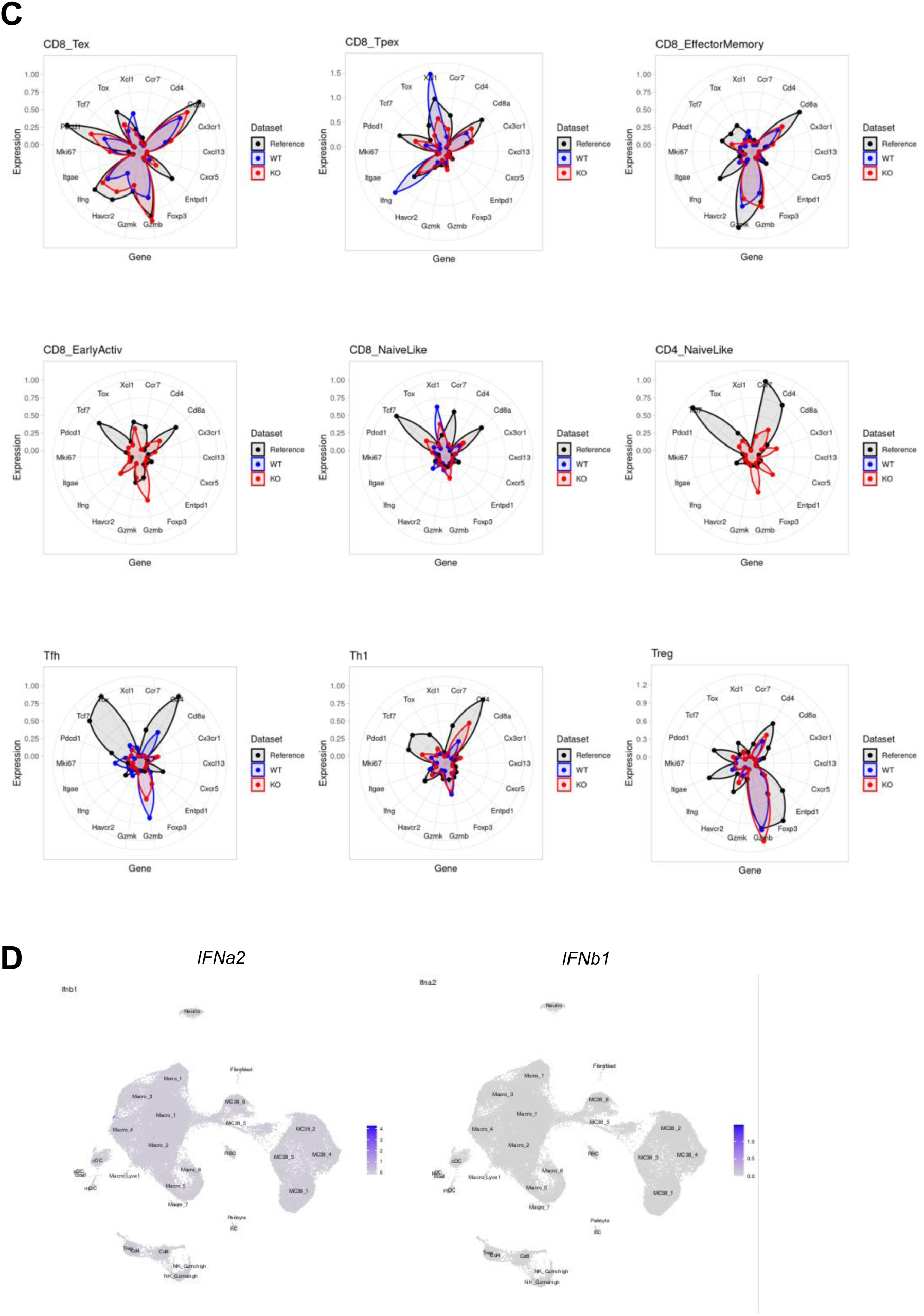
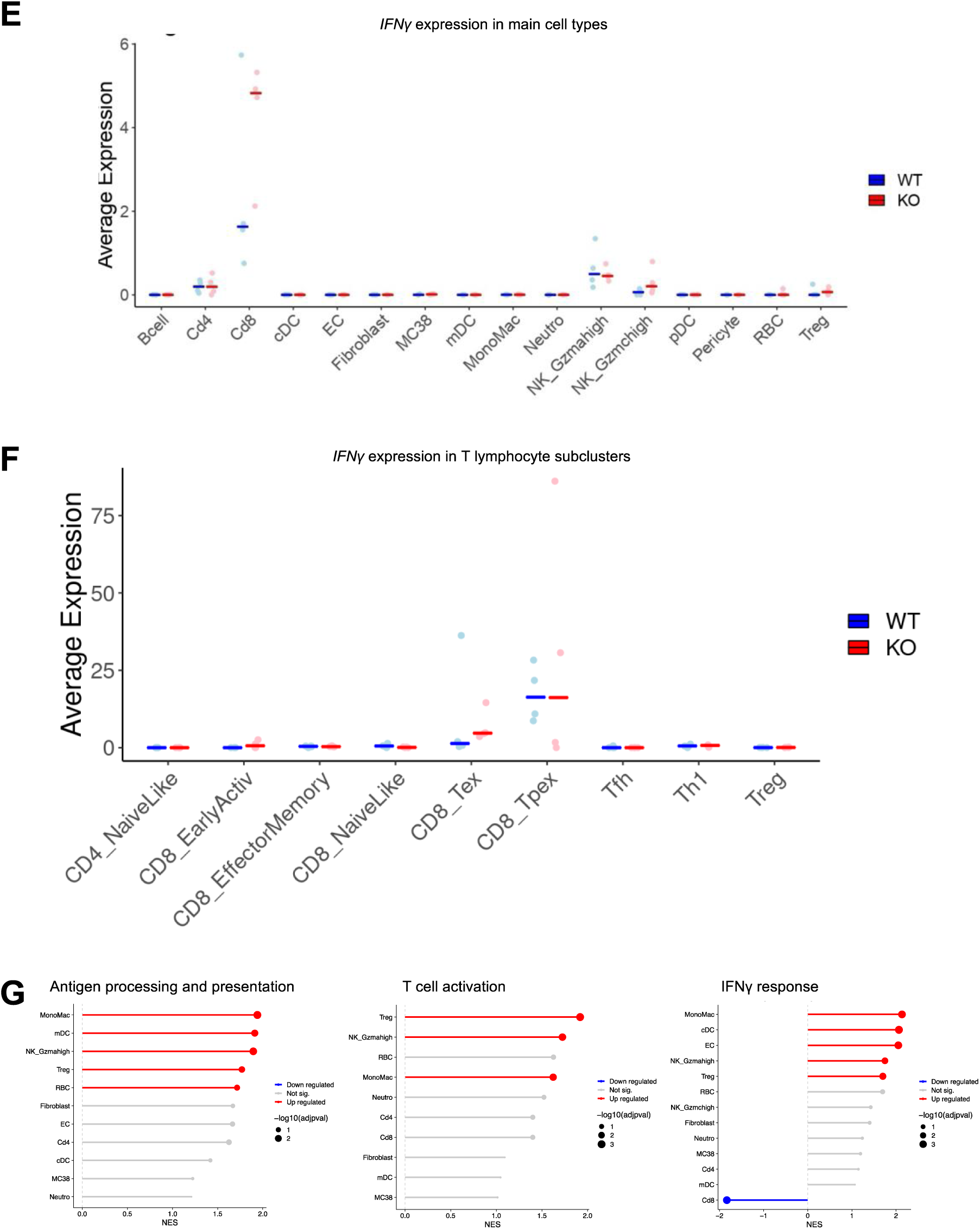
D^KO^ reprograms the microenvironment of MC38 tumors through IFNγ. A. UMAP showing the expression of selected marker genes in MC38 tumors of D^WT^ and D^KO^ mice (pooled data; *n=4* per group) by scRNA-seq. B. Bar plot showing the percentage of each cell population in MC38 tumors of D^WT^ and D^KO^ mice (pooled data; *n=4* per group). C. Radar plot representing the normalized average expression of selected genes in each T cell subset from MC38 tumors of D^WT^ and D^KO^ mice (pooled data; *n=4* per group), according to the ProjecTILs murine reference map. D. UMAP showing the expression of *Ifna2* and *Ifnb1* genes in MC38 tumors of D^WT^ and D^KO^ mice (pooled data; *n=4* per group). E. Dot plots showing the expression of the *Ifng* gene in main cell types of MC38 tumors of D^WT^ and D^KO^ mice (pooled data; *n=4* per group), Horizontal lines represent the median. F. Dot plots showing the expression of the *Ifng* gene in each T cell subset of MC38 tumors of D^WT^ and D^KO^ mice (pooled data; *n=4* per group). Horizontal lines represent the median. G. Lollipop plots showing Hallmark gene set enrichment analysis (GSEA). Genes were ranked according to log fold change between MC38 tumors of D^WT^ and D^KO^ mice (pooled data; *n=4* per group). P-values are adjusted for multiple testing according to the Benjamini-Hochberg method.

**Suppl. Fig 3.**
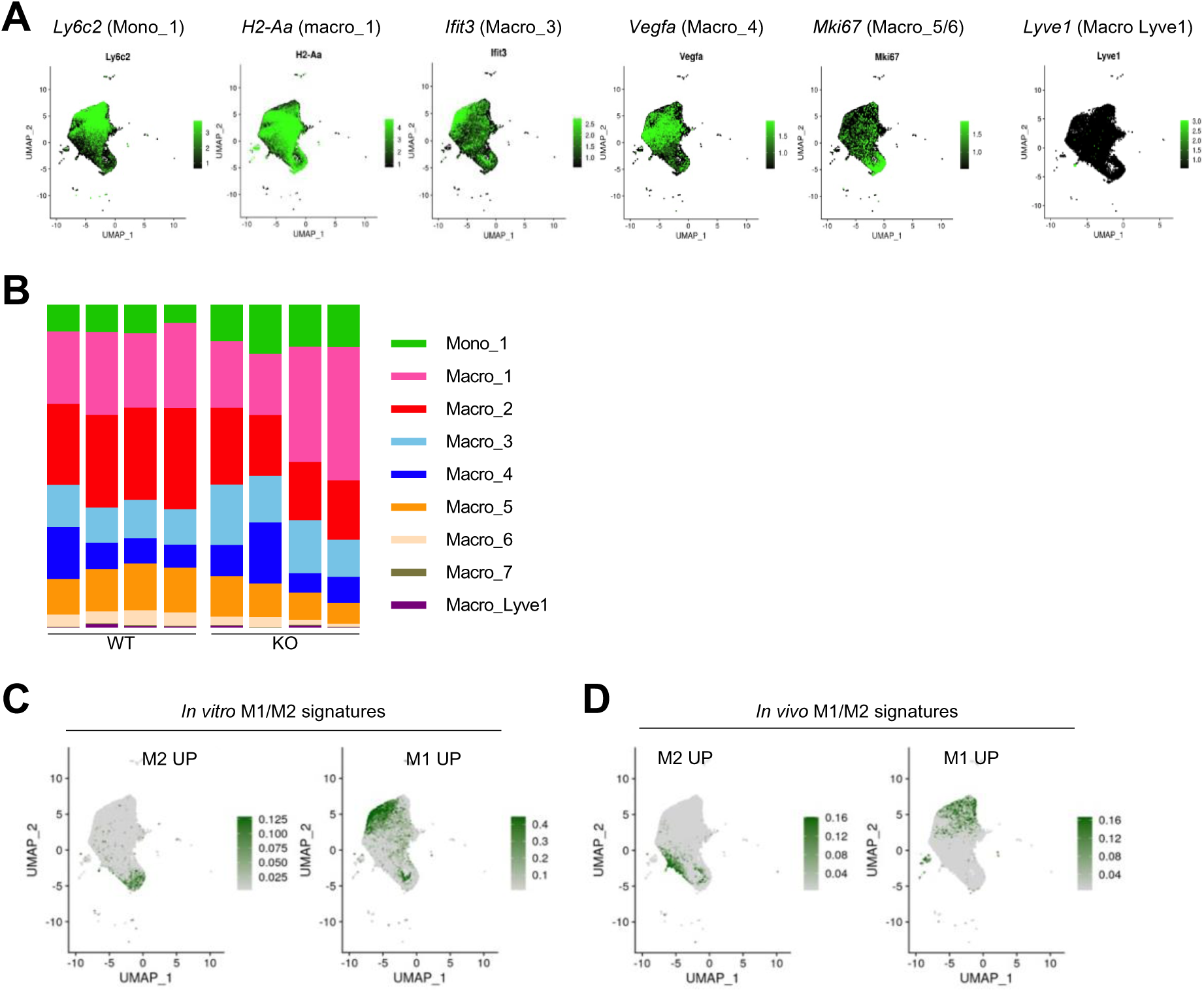
scRNA-seq reveals the co-existence of M1 and M2-like MonoMacs in MC38 tumors. A. UMAP showing the expression of selected genes in the MonoMac subclusters of MC38 tumors of D^WT^ and D^KO^ mice (pooled data; *n=4*). B. Bar plot represents the percentage of each MonoMac subcluster in MC38 tumors of D^WT^ and D^KO^ mice (*n=4*). C. UMAP showing the expression of M1/M2-like signature genes, derived from an *in vitro* dataset^28^, in MonoMac subsets of MC38 tumors of D^WT^ and D^KO^ mice (pooled data; *n=4*). D. UMAP showing the expression of M1/M2-like signature genes, derived from an *in vivo* dataset^27^, in MonoMac subsets of MC38 tumors of D^WT^ and D^KO^ mice (pooled data; *n=4*).

**Suppl. Fig 4.**
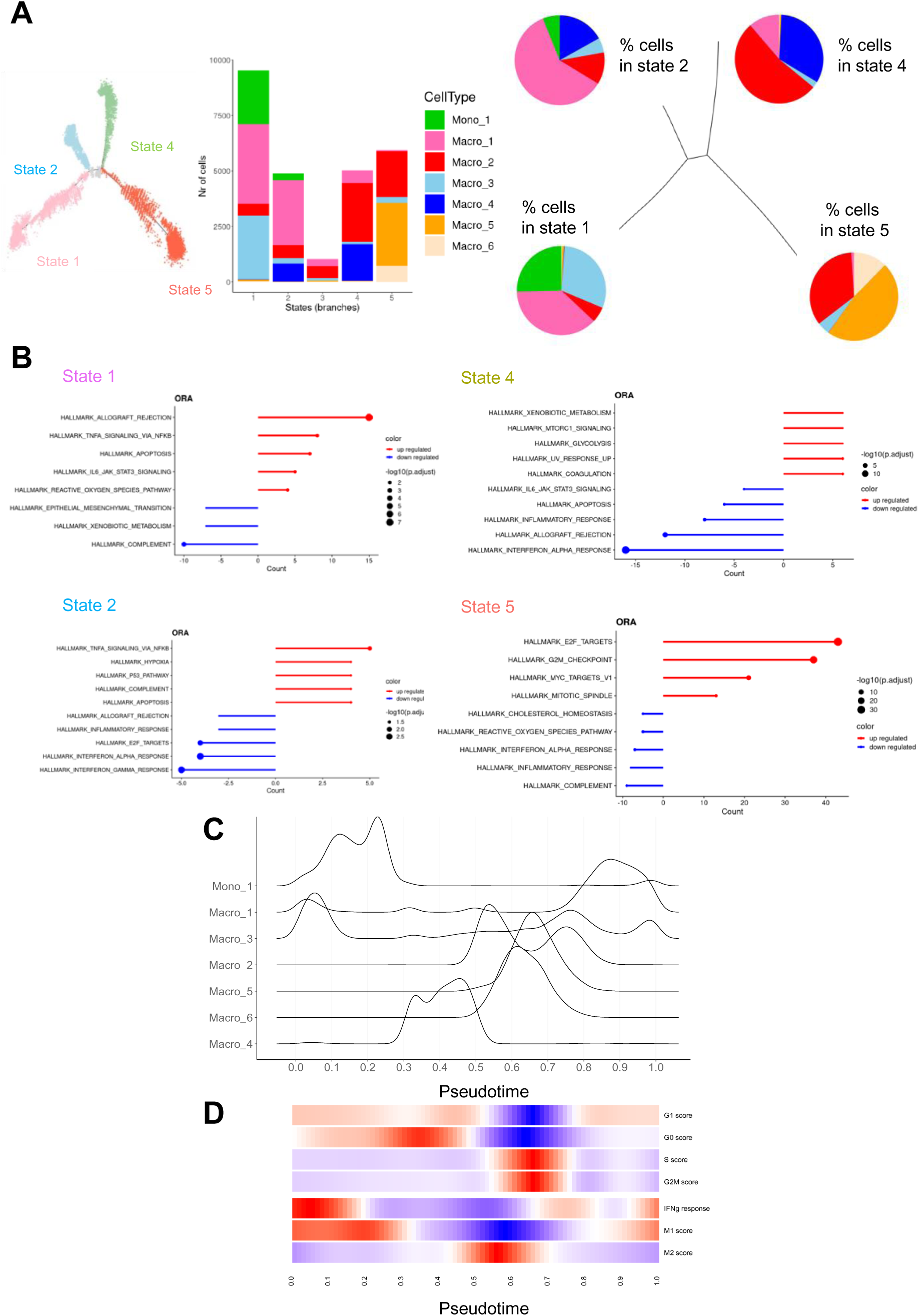
Dynamics analysis links trajectories and M1/M2 phenotypes within MonoMacsz. A. Absolute frequencies (number of cells; stacked barplot, middle) and relative proportions (percentage of cells; pie charts, right) of MonoMac subclusters across states (“branches” shown on the left) of the Monocle inferred trajectory in MC38 tumors of D^WT^ and D^KO^ mice (pooled data; *n=4*). B. Top over-(red) or under-represented (blue) Hallmark pathways in each state of the Monocle inferred trajectory. Representation (ORA) was computed from the sets of state-specific enriched genes. P-values adjusted for multiple testing according to the Benjamini-Hochberg method. C. Plot representing the density of cells in each MonoMac subcluster from MC38 tumors of D^WT^ and D^KO^ mice (pooled data; *n=4*), relative to transcriptional phase θ generated by DeepCycle. D. Heatmap showing the enrichment of cell cycle phase, M1/M2 macrophage phenotype, and Hallmark “Interferon gamma response” gene set signatures, relative to transcriptional phase θ generated by DeepCycle. M1/M2-like macrophage phenotype signatures were derived from an *in vivo* dataset^27^. Colors indicate enrichment scores, after conditional means smoothing and zero-centering.

**Suppl. Fig 5.**
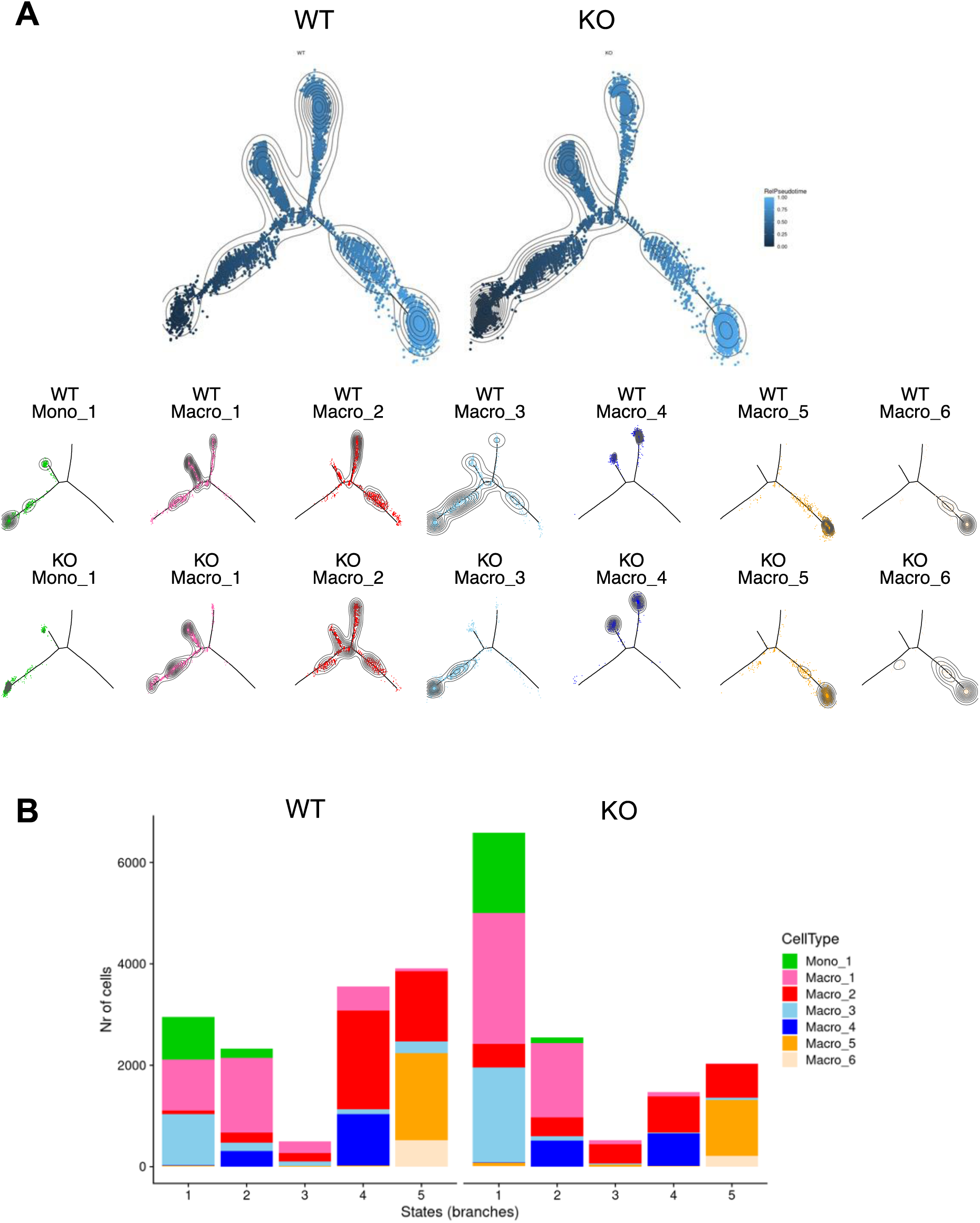
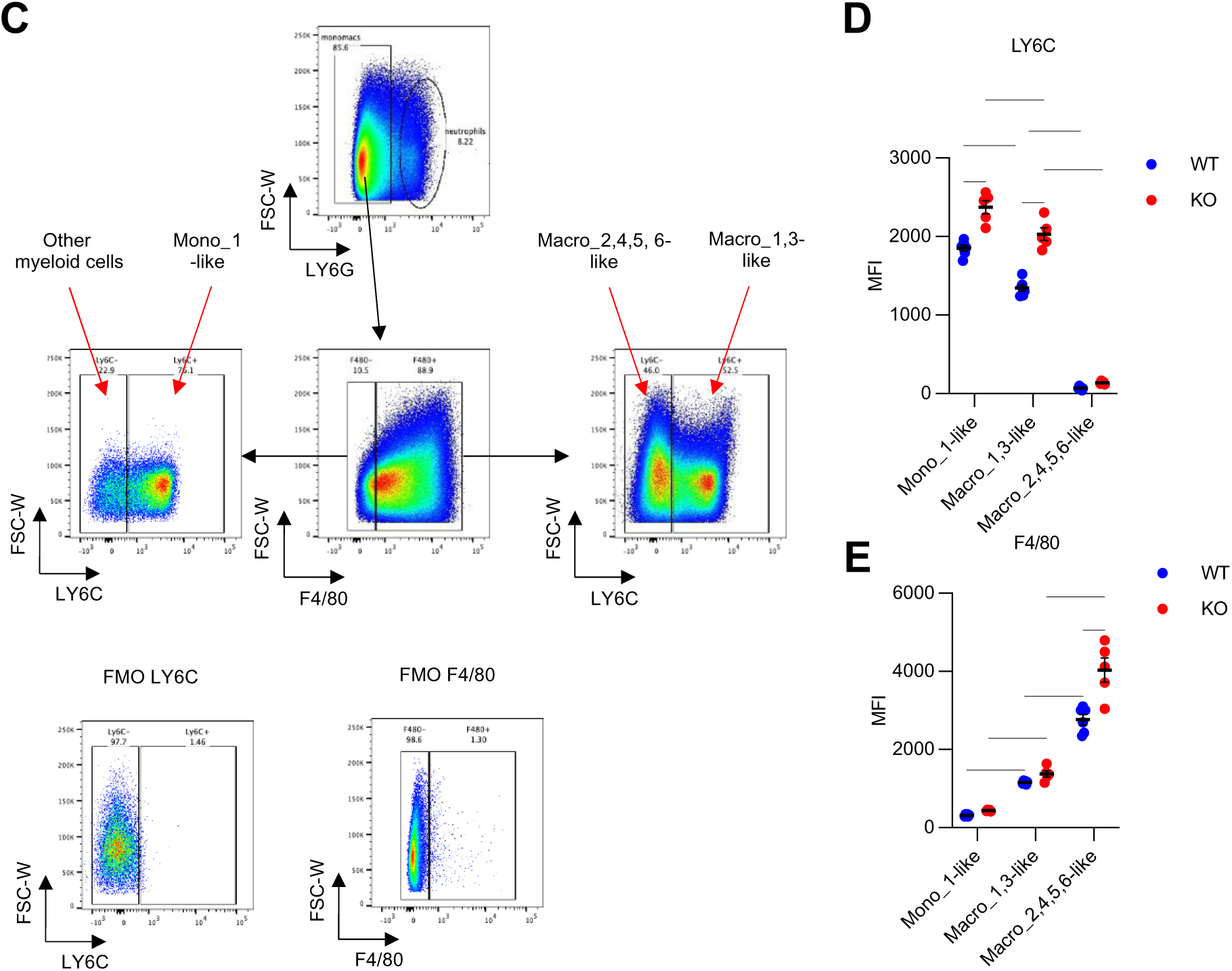
*Dicer1* inactivation in TAMs interferes with MonoMac trajectories and stalls TAMs in a cell cycle arrest state. A. Top: Distribution of relative pseudotime along the monocyte-macrophage differentiation trajectory, inferred by Monocle, in MC38 tumors of D^WT^ and D^KO^ mice (*n=4*). Concentric lines indicate the level of cell density. Bottom: Density of cells from each MonoMac subcluster across the monocyte-macrophage differentiation trajectory inferred by Monocle. Concentric lines indicate the level of cell density. B. Number of cells in the MonoMac subclusters, resolved into each state of the monocyte-macrophage differentiation trajectory inferred by Monocle, in MC38 tumors of D^WT^ and D^KO^ mice (pooled data; *n=4*). C. Representative gating strategies used to identify Macro_2,4,5, 6-like and Macro_1,3-like subsets in MC38 tumors. D. Mean fluorescent intensity (MFI) of Ly6C in each indicated myeloid subset from MC38 tumors of D^WT^ and D^KO^ mice (*n=5*). Statistical analysis by two-way Anova. E. Mean fluorescent intensity (MFI) of F4/80 in each indicated myeloid subset from MC38 tumors of D^WT^ and D^KO^ mice (*n=5*). Statistical analysis by two-way Anova.

## Methods

### Mice

C57BL/6 mice were purchased from Charles River Laboratory. For MC38 tumor studies, *Dicer1*^lox/lox^ mice (C57BL/6 background) were crossed with hemizygous C57BL/6/*LysM*-Cre mice (C57BL/6 background) to obtain *LysM*-Cre^+/WT^;*Dicer*^–/–^ mice. For the orthotopic NSCLC studies, *Dicer1*^lox/lox^ mice were crossed with homozygous C57BL/6/*LysM*-Cre mice to obtain *LysM*-Cre^+/+^;*Dicer*^–/–^ mice. Pups were genotyped by Transnetyx using qPCR assays, as detailed on the company’s website (http://www.transnetyx.com). All mice used in this study were maintained in the pathogen-free barrier animal facility of EPFL, adhering to Swiss regulations for the care and use of mice in experimental research. All procedures were performed in accordance with protocols approved by the veterinary authorities of the Canton Vaud under Swiss law (VD2916, VD3435; VD2978).

### Subcutaneous MC38 model

MC38 cells were passaged at least three times to obtain actively growing cells. The cancer cells were then resuspended in PBS at a concentration of 10 × 10^6 cells per ml, and 100 µl of the cell suspension was injected subcutaneously into the right flank of C57BL/6 mice. The tumor volume was measured using calipers, calculating the volume with the formula: tumor volume = ½ × d² × D, where D is the long diameter and d is the short diameter.

### Orthotopic NSCLC model

Confluent *Kras*^G12D^;*Tp53*^null^ NSCLC cells, established as described previously^13^, were detached using trypsin to create single-cell suspensions. The cells were then cultured in 30 μl droplets of Dulbecco’s modified Eagle’s medium with 10% fetal bovine serum for 12 hours, allowing them to aggregate. Clusters of 2000 cells were cultured in 30 μl droplets for 12 hours. Five droplets were then injected directly into the tail vein of 10-week-old C57BL/6 mice.

### Tumor monitoring by micro-CT

Micro-CT scans of the lungs of NSCLC–bearing mice were obtained using a Quantum FX micro-CT scanner (PerkinElmer). Mice were anesthetized with isoflurane throughout the imaging procedure. Imaging was performed once a week, starting from day 12 after tumor injection.

### Analysis of micro-CT scans

For measuring tumor volume (volumetry), micro-CT scans were processed and analyzed using Analyze software. 3D reconstruction was performed using Osirix-MD (Pixmeo). Tumors appeared as solid masses with a ground-glass opacity and a spherical shape. The pixel intensity range of the tumors was manually adjusted based on the contrast between the tumor and lung tissue to allow the software to automatically determine and trace the tumor margins. Manual adjustments were made when necessary. Individual tumors were defined as separate objects, and their volumes were calculated by the software.

### Tissue processing

An intracardiac perfusion with PBS was performed before organ collection.

#### Flow cytometry analysis

Tumors were minced with scissors and digested in a mixture of collagenase IV (0.2 mg/ml, Worthington), dispase (2 mg/ml, Life Technologies), and DNase I (0.1 mg/ml, Life Technologies) diluted in IMDM medium (Sigma-Aldrich) for 20 minutes at 37°C with mixing. Cell suspensions were filtered through 70 μm cell strainers and washed with FACS buffer (2% FBS, 2mM EDTA in PBS) before antibody staining.

#### qPCR analysis

Tumors were harvested, snap-frozen in liquid nitrogen, and stored at –80°C.

#### Immunofluorescence staining of lung tumors

Tumors were dissected from the lung and embedded in O.C.T. compound (Cryomatrix, Thermo Fisher Scientific) and subsequently placed on dry ice. Blocks were stored at – 80°C. Tissue sections of 8 μm were obtained using a Leica cryostat CM1950 (Leica Biosystems) and stored at –20°C.

### Immunofluorescence staining of lung sections

For CD4 and CD8 staining, lung tumor sections were first fixed in methanol for 20 minutes at –20°C, washed three times with PBS for 5 minutes each, and incubated with blocking solution containing 1% bovine serum albumin (BSA) and 5% FBS in PBS for 1 hour at room temperature. Sections were then incubated overnight at 4°C in 100-200 μL of blocking solution containing Rat anti-CD8α-AF647 (1:50, clone 53-6.7, 557682, BD Bioscience) and Rat anti-CD4-PE (1:100, clone RM4-5, 100512, Biolegend). After staining, nuclei were labeled with DAPI (1 μg/ml, Sigma Aldrich) and sections mounted in Dako fluorescence mounting medium, covered with cover glass (Heathrow Scientific), and stored at 4°C. Slides were washed three times with PBS for 5 minutes between each step. Images were acquired using Zeiss LSM700 confocal microscope to acquire immunofluorescent images of tumors that were previously identified on adjacent hematoxylin/eosin-stained sections. Images were analyzed with Image J without knowledge of the treatment group. A region of interest (ROI) of the tumor area was defined before any parameter analysis.

### Flow cytometry

Tumor cell suspensions were incubated with Fc block (1:100, clone 2.4G2, 553142, BD Bioscience) before surface marker staining with combinations of the following antibodies:

CD45 APC-eF780 (1:200 clone 30-F11, 47-0451-82, ebioscience)

CD3-BV605 (1:200, clone 17A2, 100237, Biolegend)

CD4-PE (1:200, clone RM4-5, 100512, Biolegend)

CD4-PB (1:200, clone RM4-5, 100547, Biolegend)

CD8α-PE (1:50, clone YTS169.4, MA182111, ThermoFisher Scientific)

CD11b-BV711 (1:200, clone M1/70, 101241, Biolegend)

MRC1-AF647 (1:100, clone C068C2, 141717, Biolegend)

Ly6C-AF700 (1:100, clone HK1.4, 128024, Biolegend)

Ly6G-PerCP Cy5.5 (1:200, 1A8, 127615, Biolegend)

F4/80-AlexaFluor488 (1:100, clone BM8, 123119, Biolegend) LIVE/DEAD Dapi for dead cell discrimination (Thermo Fisher)

Cell staining was performed in darkness at 4°C for 20-30 minutes in FACS buffer, except for LIVE/DEAD staining, which was performed in PBS. Cells were washed between each step with FACS buffer, PBS, and resuspended in FACS buffer before analysis with an LSRII SORP apparatus (BD Biosciences). Data were analyzed using FlowJo. Cells were gated based on their size and granularity (FSC-A versus SSC-A), followed by doublet and dead-cell exclusion.

The distinct CD45^+^ hematopoietic cell types were identified as follows:

Tumor-associated macrophages (TAMs): CD11b^+^ Ly6G^−^ F4/80^+^ MRC1^+^

Monocytes: Ly6G^−^ F4/80^−^ CD11b^+^ Ly6C^+^

Neutrophils: CD11b^+^ Ly6G^+^

CD3^+^ T cells: CD11b^−^ Ly6G^−^ F4/80^−^ CD3^+^

CD4^+^ T cells: CD11b^−^ Ly6G^−^ F4/80^−^ CD3^+^ CD4^+^

CD8^+^ T cells: CD11b^−^ Ly6G^−^ F4/80^−^ CD3^+^ CD8^+^

CD11b^+^ cells: CD11b^+^ Ly6G^−^ F4/80^+^

### Treatment of mice with neutralizing antibodies

The IFNγ-neutralizing antibody (12 mg kg^–1^; clone XMG1.2, rat IgG1, Bio X Cell) was administered intraperitoneally twice per week starting from day 7 after MC38 tumor injection, until the end of the experiment. The Anti-VEGFA/ANGPT2 (20 mg/kg; murinized A2V IgG2a, Roche) and anti–PD-1 (10 mg/kg; rat IgG2a, clone RMPI-14, BE0146, Bio X Cell) antibodies were administered on days 15, 22 and 29 post-NSCLC injection. For the latter study, irrelevant IgGs were mouse IgG1 (20 to 30 mg/kg; clone MOPC-21, Roche) for A2V, and rat IgG2a (10 mg/kg; clone 2A3, BE0089, Bio X Cell) for anti–PD-1.

### IFNγ measurement using qPCR and ELISA

For RNA extraction and qPCR analysis, total mRNA was extracted using the RNeasy Mini Kit (Qiagen). Briefly, samples were homogenized in 700 μl of QIAzol lysis reagent, followed by addition of chloroform and centrifugation. The upper phase was transferred to a new tube, mixed with ethanol, and loaded onto RNeasy Mini columns. After washing steps with Buffer RWT and Buffer RPE, RNA was eluted in RNase-free water. RNA concentration was quantified using NanoDrop ND-2000. cDNA was synthesized using the SuperScript Vilo cDNA Synthesis Kit (Invitrogen). RNA (up to 20 μl) was mixed with 4 μl of 5x VILO Reaction Mix and 2 μl of 10x SuperScript Enzyme Mix. Reverse transcription was performed at 25°C for 10 min, 42°C for 60 min, and 85°C for 5 min. cDNA samples were stored at -20°C. TaqMan gene assays (Applied Biosystems/Life Technologies, *Ifng* probel Mm01168134_m1) were used. Each reaction contained 10 ng of cDNA per well in triplicate in a 384-well plate. A master mix was prepared with qPCR Universal master mix and TaqMan gene assay. qPCR was run for 40 cycles on an ABI7900HT apparatus. Raw data were analyzed using SDS software v2.4. Relative gene expression levels were calculated using the ΔΔCt method normalized to a reference gene (*Gapdh*).

Serum IFNγ protein was measured using the mouse IFNγ ELISA kit (StemCell Technologies 02020), following the manufacturer’s instructions.

### Statistical analysis

Graphs were generated and statistical analyses were performed using Prism (GraphPad Software). Error bars represent the standard error of the mean (SEM), unless indicated otherwise. The number of biological (nontechnical) replicates and the specific statistical analyses used are detailed in the figure legends. Data distribution normality was tested using the Shapiro-Wilk normality test. Outliers were not excluded from the analysis. Comparisons between two unpaired groups were performed using the parametric Student’s *t*-test. For multiple comparisons, two-way ANOVA was used. P values are indicated as *: P ≤ 0.05, **: P ≤ 0.01, ***: P ≤ 0.001, and ****: P ≤ 0.0001 in all figures.

### Single cell RNA sequencing

#### Sample preparation

We conducted single-cell RNA sequencing (scRNA-seq) of MC38 tumors obtained from either *Dicer*-WT or *Dicer*-KO mice at days 11-12 after MC38 tumor challenge. Tumors were minced with scissors and digested in a mixture of collagenase IV (0.2 mg/ml, Worthington), dispase (2 mg/ml, Life Technologies), and DNase I (0.1 mg/ml, Life Technologies) diluted in IMDM medium (Sigma-Aldrich) for 20 minutes at 37°C with mixing. Single cells were filtrated through a 40 μm Flowmi strainer (Bel-Art) and resuspended in PBS with 0.04% BSA, checked for the absence of doublets or aggregates and loaded into a Chromium Single Cell Controller (10× Genomics, Pleasanton) in a chip together with beads, master mix reagents (containing reverse transcriptase and poly-dT primers) and oil to generate single-cell-containing droplets.

#### Data acquisition

Single-cell gene expression libraries were prepared using Chromium Single Cell 3’ Library & Gel Bead Kit v2 following the manufacturer’s instruction. Quality control was performed with a TapeStation 4200 (Agilent) and QuBit dsDNA high sensitivity assay (Thermo fisher scientific) following manufacturer instructions. The libraries were sequenced on an Illumina HiSeq4000 platform, with run conditions as per 10X recommendations, aiming at 50’000 reads/cell.

#### Data processing

The Cell Ranger Single Cell Software Suite v3.1.0 was used to perform sample demultiplexing, barcode processing, and 3’ gene counting using 10X Genomics custom annotation of murine genome assembly mm10-3.0.0. Data was analyzed using the standard functions from the Seurat R package (v. 4.0.4). Breifly, cells above 1% and below 99% of total feature counts were kept. Cells with more than 10% of features mapping to the mitochondrial genome were discarded. Raw data was integrated, scaled and log-transformed; principal component analysis (PCA) was performed based on the expression of the top 2000 most variable genes, selected using the variance-stabilizing transformation (“vst”) method; Uniform Manifold Approximation and Projection (UMAP) dimensional reduction was performed based on the first 20 principal components. Unsupervised clustering was performed applying the graph-based clustering approach and Louvain algorithm at different resolutions, based previously computed neighbor graph using the top 30 PCs. Clusters were identified and merged into cell populations based on cell type specific markers.

#### Differential and pathway enrichement analyses

Differential gene expression was computed within each cell population between conditions using the pseudo-bulk approach implemented in muscat (v1.16.0), using the method edgeR, or using FindAllMarkers. Genes expressed in at least 10% of the cells per population per sample and with average logCPM > 5 were retained. Gene set enrichment analysis was performed with clusterProfiler (v.4.0.5) applying GSEA with default parameters and 100’000 permutations to obtain P-values. Overrepresentation analysis was performed on a subset of regulated genes applying clusterProfiler enrichr. The Hallmark collection from msigdbr v.7.4.1 was used, or specific gene signatures extracted from published studies or in mSigDB. Differential abundance of cell populations between D^WT^ and D^KO^ mice was based on Student’s *t-*test for comparison of means. P-values were adjusted for multiple testing using the Benjamini-Hochberg method.

#### Prediction of T lymphocyte cellular states

The R package ProjecTILs (v 2.0.3) was used to project T lymphocyte transcriptomic profiles onto a reference map of tumor-infiltrating lymphocytes (TILs) cellular states ^15^. The pre-computed cross-study pan-cancer murine TIL Atlas version 1.0 (http://tilatlas.unil.ch/) was downloaded and used as the reference map. Prediction was performed using a nearest-neighbor classifier and was based on a majority vote of its annotated nearest neighbors in PCA space. The first ten principal components were used to compute distances, and the majority vote was based on the 20th nearest neighbors.

#### Cell cycle phase enrichment scores

G1, S and G2M cell cycle phases were scored using the CellCycleScoring function from the Seurat R package (v. 4.0.4) ^45^, and the cell cycle phase markers from Tirosh *et al.* ^36^. The G0 cell cycle phase signature was obtained from Rappez *et al.* ^37^ and scored using the AddModuleScore_Ucell ^46^ function from the Seurat R package (v. 4.0.4), which computes the Mann-Whitney U statistic implemented in the Ucell R package (v. 1.3.1).

#### M1/M2-like signatures and IFNγ response enrichment scores

Tumor-associated macrophage signatures (M1-like and M2-like) were obtained from references ^27,28^. The signature for IFNγ response consisted of the Hallmark pathway “Interferon Gamma response” gene set, and was obtained from the Molecular Signatures Database (MsigDB) ^47–49^ through the msigdbr R package (v. 7.4.1) https://igordot.github.io/msigdbr/. Signature enrichments were scored using the AddModuleScore_Ucell^46^ function from the Seurat R package (v. 4.0.4). ***Trajectory analysis of MonoMac sub-populations*.** Trajectory analysis of MonoMac sub-populations was performed using functions from the Monocle R package (v. 2.20.0) ^33–35^. Only cells expressing at least 500 genes, and genes expressed in at least 25% of cells that were also positive markers for the MonoMac sub-populations, were considered. Markers were identified using the FindAllMarkers ^45^ function from the Seurat R package, using a threshold for the minimum average log2 fold change of 0.35. Differential expression was tested using the hurdle model implemented in the MAST R package (v. 1.18.0) https://github.com/RGLab/MAST/. The trajectory was learned by first reducing the dimensionality of the data (reduceDimension function) and then ordering cells according to a pseudotime (orderCells function). Dimensionality was reduced to two using the Discriminative Dimensionality Reduction with Trees algorithm implemented in the DDRTree R package (v. 0.1.5) https://github.com/cole-trapnell-lab/DDRTree. Cells were ordered without any pre-determined “root” state or number of end-point cell states. Positive markers for different states in the trajectory were determined using the *FindAllMarkers* function from Seurat, ranked according to log fold change, and used for performing over-representation analysis (ORA) of Hallmark pathways gene sets (obtained through the msigdbr R package). ORA was performed using the *enricher* function from the clusterProfiler R package (v. 4.0.5)^50^.

#### Assignment of a continuous cell cycle trajectory to MonoMac cells

Cell cycle trajectory analysis of MonoMac cells was performed using DeepCycle ^51^. First, we used Velocyto (v. 0.17) ^52^ to create loom files with read counts divided by spliced, unspliced or ambiguous mRNA. Loom files were then used to compute the moments for RNA velocity estimation using scVelo (v.0.2.4) ^53^. Finally, the results from RNA velocity were used to run Deepcycle in two steps. In the first step, a random gene from the provided list of mouse cycling genes (GOterm:cell_cycle, GO:0007049) was used as the initial condition for the transcriptional phase, together with the hotelling filter and an expression threshold of 0.1. In the second step, Actb was chosen as the gene for the initial condition for the transcriptional phase, the hotelling filter was not used, and the expression threshold was set to 0.5.

#### Cell cycle phase enrichment scores

In Fig 4F, G1, S and G2M cell cycle phases were scored using the CellCycleScoring ^45^ function from the Seurat R package (v. 4.0.4), and the cell cycle phase markers from Tirosh et al. 2015 ^36^. The G0 cell cycle phase signature was obtained from Rappez et al. 2020 ^37^ and scored using the AddModuleScore_Ucell ^46^ function.

#### M1/M2-like signatures and IFNγ response enrichment scores

In Fig 4F, tumor-associated macrophage signatures (M1-like and M2-like) were obtained from the following references: Refs ^27,28^. The signature for IFNγ response consisted of the Hallmark pathway “Interferon Gamma response” gene set, and was obtained from the Molecular Signatures Database (MsigDB) through the msigdbr R package ^48^. Signature enrichments were scored using the AddModuleScore_Ucell^46^ function.

#### Differential abundance of MonoMac subsets

The analysis of differential abundance of cell populations between D^WT^ and D^KO^ mice was based on Student’s *t*-test for comparison of means. P-values were adjusted for multiple testing using the Benjamini-Hochberg method.

### Macrophage signatures in human cancer

The hepatocellular carcinoma (HCC) and melanoma (MEL) human single-cell RNA sequencing data were obtained from Gene Expression Omnibus (GEO: GSE154763) ^42^. Data were analyzed using the standard functions from the Seurat R package (v. 4.0.4). Raw data was log-normalized, scaled and centered. Principal component analysis (PCA) was performed based on the expression of the top 2000 most variable genes, selected using the variance-stabilizing transformation (“vst”) method. Uniform Manifold Approximation and Projection (UMAP) dimensional reduction was performed based on the first ten principal components. Cell cycle phases and tumor associated macrophage signatures were scored following the procedure described above.

### Data Availability

The scRNA sequencing data generated in this study have been deposited in the Gene Expression Omnibus (GEO) under accession number GSE274402.

